# Acute aerobic exercise intensity does not modulate pain potentially due to differences in fitness levels and sex effects – results from a pharmacological fMRI study

**DOI:** 10.1101/2024.06.25.600579

**Authors:** Janne I. Nold, Tahmine Fadai, Christian Büchel

## Abstract

Exercise might lead to a release of endogenous opioids, potentially resulting in pain relief. However, the neurobiological underpinnings of this effect remain unclear. Using a pharmacological within-subject fMRI study with the opioid antagonist naloxone and different levels of aerobic exercise and pain we investigated exercise-induced hypoalgesia (*N* = 39, 21 female). Overall, high-intensity aerobic exercise did not reduce pain as compared to low-intensity aerobic exercise. Accordingly, we observed no significant changes in the descending pain modulatory system. The µ-opioid antagonist naloxone significantly increased overall pain ratings but showed no interaction with exercise intensity. An exploratory analysis suggested an influence of fitness level (as indicated by the functional threshold power) and sex where males showed greater hypoalgesia after high-intensity exercise with increasing fitness levels. This effect was attenuated by naloxone and mirrored by fMRI signal changes in the medial frontal cortex, where activation also varied with fitness level and sex, and was reversed by naloxone. These results indicate that different aerobic exercise intensities have no differential effect on pain in a mixed population sample, but individual factors such as fitness level and sex might play a role. The current study underscores the need for personalised exercise interventions to enhance pain relief in healthy as well as chronic pain populations taking into account the sex and fitness status as well as the necessity to further investigate the opioidergic involvement in exercise-induced pain modulation.

## Introduction

Exercise has been suggested to release endogenous opioids (Koltyn, 2000) resulting in an attenuation of pain, termed exercise-induced hypoalgesia. However, the underlying neural (Vaegter and Jones, 2020; Wu et al., 2022) and pharmacological mechanisms remain controversial (Koltyn, 2000; Naugle et al., 2012; Wewege and Jones, 2020) and results on the modulation of pain through exercise are heterogeneous (Klich et al., 2018; Koltyn et al., 2014; Padawer and Levine, 1992; Ruble et al., 2005; Sternberg et al., 2001; Vaegter et al., 2015, 2019). Given that exercise can potentially elicit hypo- and hyperalgesia (Vaegter and Jones, 2020) we have selected the term ‘exercise-induced pain modulation’ to reflect the range of effects of exercise on pain.

Previous rodent research has implicated the descending pain modulatory system, including the anterior cingulate cortex (ACC), the medial frontal cortex including the ventromedial prefrontal cortex (vmPFC), and the periaqueductal grey (PAG) in exercise-induced pain modulation (Lesnak and Sluka, 2020; Stagg et al., 2011). This is partly congruent with early imaging studies in humans (Boecker et al., 2008; Ellingson et al., 2016; Geisler et al., 2019; Saanijoki et al., 2018; Scheef et al., 2012), where the descending pain modulatory system, including the PAG and insula, seems to be involved in modulating pain perception following a 2-h run (Scheef et al., 2012) along with the fusiform gyrus and hippocampus (Geisler et al., 2019).

The activation of the endogenous opioid system during exercise has been the most widely discussed mechanism in exercise-induced pain modulation (Naugle et al., 2012; Sluka et al., 2018). Studies frequently employ the µ-opioid antagonist naloxone which blocks opioid receptors in the brain, thus counteracting opioid-driven mechanisms of pain modulation. Earlier studies found an analgesic response to thermal and ischaemic pain after long-distance (6.3 miles) running in male runners that was reversed after naloxone administration (Janal et al., 1984). Even after a short-distance (1 mile) run, naloxone slowed the return of the analgesic effect (Haier et al., 1981). Previous imaging studies (Boecker et al., 2008; Saanijoki et al., 2018) further supported that exercise elicits the release of endogenous opioids in crucial opioidergic structures (i.e., the insula, ACC, and prefrontal regions). Still, the pharmacological underpinnings in humans have remained equivocal (Koltyn, 2000), where administering a µ-opioid antagonist in some studies decreased exercise-induced pain modulation (Droste et al., 1988; Haier et al., 1981; Janal et al., 1984; Olausson et al., 1986) whereas in others it did not affect exercise-induced pain modulation (Droste et al., 1991; Koltyn et al., 2014; Olausson et al., 1986). Notably, most studies investigating the pharmacological underpinnings of exercise-induced pain modulation have employed very different interventions using different administration protocols (i.e., different doses and different routes of administration of the µ-opioid receptor antagonist such as oral or intravenous), making it difficult to draw clear conclusions about the extent of central µ-opioid modulation through exercise in humans (Koltyn, 2000; Leknes and Atlas, 2020; Wewege and Jones, 2020).

In human studies, many findings on exercise-induced pain modulation are derived from studies in athletes or physically active individuals showing increased hypoalgesia following exercise compared to subjects with lower fitness levels (Crombie et al., 2018; Geva and Defrin, 2013; Naugle and Riley, 2014; Schmitt et al., 2020). Intriguingly, this observation might be related to greater exercise-induced opioid release in structures such as the PAG or the frontal medial cortex (including the ACC) in fitter individuals (Saanijoki et al., 2022; Sluka et al., 2018). Additionally, previous reviews have further emphasised sex differences in pain processing (Mogil, 2020, 2012) and analgesic sensitivity in rodents (Martin et al., 2019; Mogil, 2020) warranting further investigation in humans (Mogil, 2020, 2012) in connection with exercise-induced pain modulation.

Furthermore, studies on exercise-induced pain modulation have been criticised (Padawer and Levine, 1992) for using single-arm pre-post measurements where pain is measured once before and once after the exercise intervention instead of randomised trials with the former not accounting for habituation to the noxious stimulus (for reviews see Naugle et al., 2012; Vaegter and Jones, 2020), as well as heterogeneous study designs (Vaegter and Jones, 2020), further limiting the interpretation of results (Naugle et al., 2012; Wewege and Jones, 2020).

In this study, we examined exercise-induced pain modulation between different exercise intensities in a healthy sample with a particular emphasis on the opioidergic system. This preregistered study (drks.de: DRKS00029064) comprised an initial calibration day and two experimental days (Figure 1). On the calibration day, individual exercise intensities, pain intensities, as well as objectively assessed fitness levels (functional threshold power; FTP_20_) (Borszcz et al., 2018), were determined (Figure 1A). On each of the two experimental days, participants cycled during four blocks at high (100% FTP) and low (55% FTP; active recovery) intensity for 10 minutes per block. After each cycling block, functional Magnetic Resonance Imaging (fMRI) commenced, where participants received nine painful heat stimuli (alternating with nine painful pressure stimuli) with a 15-second plateau each and rated their painfulness on a visual analog scale (VAS) ranging between the calibrated pain threshold (‘minimally painful’) and pain tolerance (‘almost unbearably painful’). Furthermore, we employed a double-blind cross-over pharmacological challenge across experimental days, where either saline (placebo; SAL) or the µ-opioid antagonist naloxone (NLX) was administered intravenously before the experiment commenced and a constant dose was maintained throughout the experiment (see Methods for details) (Figure 1B).

**Figure 1.**
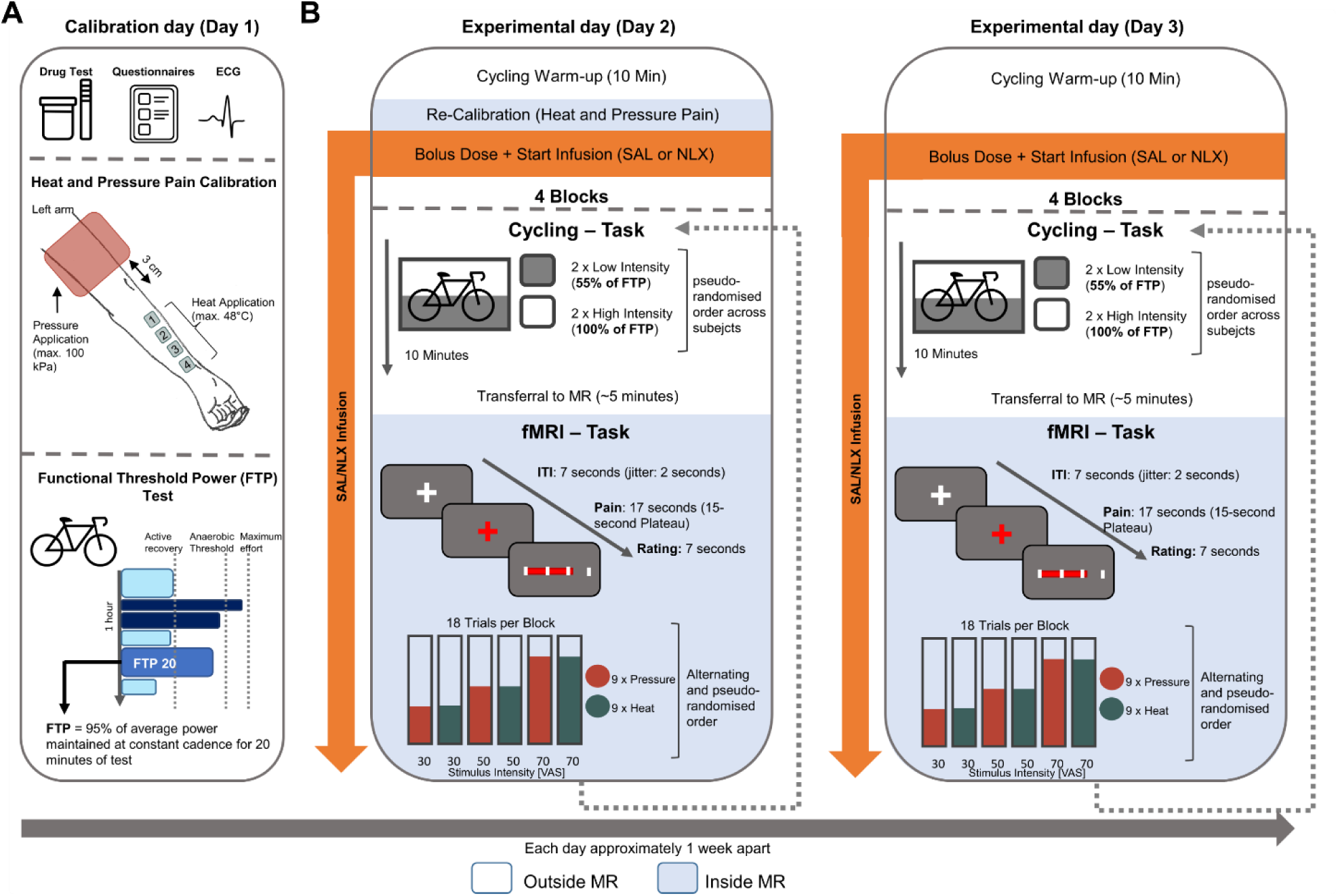
Experimental design. (**A**) Calibration day (Day 1) with heat and pressure pain calibration and Functional Threshold Power test (FTP) (see Table S1). (**B**) Experimental days (Day 2 and Day 3) with cycling task (outside MR) and fMRI task (inside MR) with the only difference in the drug treatment administered (SAL or NLX). ITI = inter-trial-interval; VAS = Visual Analog Scale.

## Results

In our preregistration, we also included pressure pain to investigate exercise-induced pain modulation across different pain modalities. However, pressure pain did not show a significant effect of exercise in any of the analyses performed (Figure S1); therefore, this report focuses on heat pain. As the inter-trial interval (ITI) between heat pain stimuli needs to be about 30 seconds due to the stimulus length (Tran et al., 2010), the interspersed pressure pain stimuli allowed us to additionally (i) characterise the brain responses to pressure as compared to heat pain and (ii) investigate the effect of a µ-opioid antagonist on these responses. However, these results are beyond the scope of this paper and will be reported separately. The final sample included healthy female (*n* = 21) and male (*n* = 18) participants of varying fitness levels. The equal distribution of menstrual phases (based on self-report and divided into three phases: follicular, ovulatory, and luteal) for the experimental days was estimated using a χ^2^ -test (Wilson, 1927) and showed neither significant differences on each experimental day nor between days (see Table S2).

### Effective induction of heat pain

As a first step, we verified the successful application of the calibrated heat pain intensities in the placebo (saline; SAL) condition. A significant main effect of stimulus intensity on heat pain ratings was observed (*β* = 1.42, *CI* [1.37,1.48], *SE* = 0.03, *t*(1358.65)= 51.84, *P* < 2×10^−16^; Figure 2A, Table S3, and S4). Corroborating this, our fMRI data showed blood oxygenation level-dependent (BOLD) activation changes in key brain regions associated with pain processing such as the right anterior Insula (antIns; Figure 2B; MNI_xyz_: 36, 6, 14, T = 10.35, *P_corr-WB_* < 0.001), right dorsal posterior Insula (dpIns; Figure 2C; MNI_xyz_: 39, - 15, 18, T = 7.65, *P_corr-WB_* < 0.001) and right middle cingulate cortex (MCC; Figure 2D; MNI_xyz_: 6, 10, 39, T = 7.47, *P_corr-WB_* = 0.001). The parameter estimates from the respective peak voxels mirrored the behavioural response pattern (Figure 2E-G). Overall, this confirms the successful administration of thermal heat stimuli at the intended intensities. For an uncorrected activation map at *P_uncorr_* < 0.001 of the entire brain see Figure S2.

**Figure 2.**
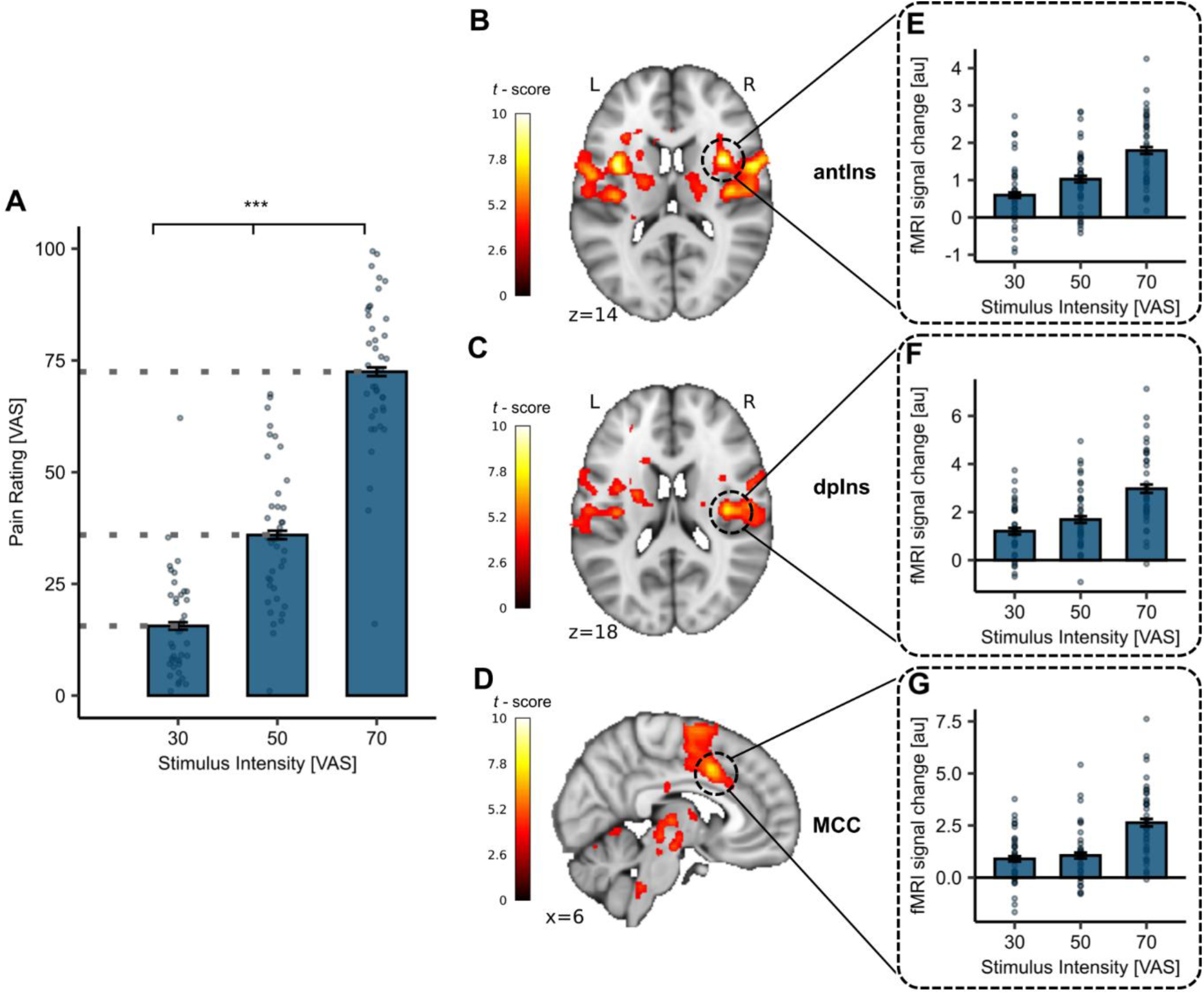
Behavioural and fMRI results for successful pain induction. (**A**) Heat pain ratings in the SAL condition for all stimulus intensities (VAS 30, 50, 70) showed a significant main effect of stimulus intensity (*P* < 2×10^−16^). The *P*-value reflects the significant main effect of stimulus intensity from the linear mixed effect model (LMER). (**B-D**) Significant activation for the parametric effect (Heat VAS 70 > 50 > 30) in the SAL condition in the (B) right antIns (MNI_xyz_: 36, 6, 14; T = 10.35, *P_corr-WB_* < 0.001), (C) right dpIns (MNI_xyz_: 39, −15, 18, T = 7.65, *P_corr-WB_* < 0.001) and (D) right MCC (MNI_xyz_: 6, 10, 39; T = 7.47, *P_corr-WB_* = 0.001). Displayed are the uncorrected activation maps (*P_uncorr_* < 0.001) for visualisation purposes. (**E-G**) Bars depict mean parameter estimates from the respective peak voxels for all stimulus intensities across participants whereas dots display subject-specific mean parameter estimates at the respective stimulus intensity. *P*-values were calculated using the LMER model for the fixed effect of stimulus intensity. * *P* < 0.05, **, *P* < 0.01, *** *P* < 0.001. Error bars depict SEM (*N* = 39).

### Implementation of exercise intervention

As a next step, we aimed to confirm the successful implementation of our exercise intervention. This intervention comprised previously calibrated high-intensity (HI) and low-intensity (LI) exercise conditions, where the anticipated intensities were derived from the individual functional threshold power (FTP) value determined on Day 1 (Figure 1A). The anticipated power during HI power was set to 100% whereas LI power was set to 55% of FTP. As intended, the maintained power during cycling was significantly higher in the HI exercise condition compared to the LI exercise condition (*t*(37) = 16.09, *P* < 2.2×10^−16^, mean power_diff_ = 60.26W, *d* = 0.61, *n* = 38). Furthermore, the relative Power (as % of FTP) was significantly different between the exercise conditions (*t*(37) = 44.58, *P* < 2.2×10^−16^, *d* = 6.46, *n* = 38; Figure 3A). Supporting this, the mean absolute heart rate was significantly higher in the HI exercise condition compared to the LI exercise condition (*t*(33) = 20.48, *P* < 2.2×10^−16^, mean bpm_diff_ = 26.92, *d* = 1.85, *n* = 34, Figure 3B). In one participant the power recording was incomplete (*n* = 38) and five participants had faulty HR recordings in one session (*n* = 34) due to an instrumentation error. Nevertheless, the actual power and HR maintained throughout the blocks were also visually monitored. Furthermore, participants rated their perceived level of exertion (RPE) on a BORG scale (Borg, 1998) from 6 to 20 following each exercise block. The mean RPE across all HI and LI exercise blocks was also significantly higher after HI compared to LI exercise (*t*(38) = 19.65, *P* < 2.2×10^−16^, mean RPE_diff_ = 6.32, *d* = 3.69, *n* = 39; Figure 3C). Thus, the calibration and implementation of the exercise intensities were successful. To account for a potential expectation about the effect of acute exercise on pain we have conducted a questionnaire (Lindheimer et al., 2020) on the calibration day (Day 1) to capture this. However, there were no significant expectation effects on different pain domains evident (Figure S3) suggesting that expectation is unlikely to have influenced the results.

**Figure 3.**
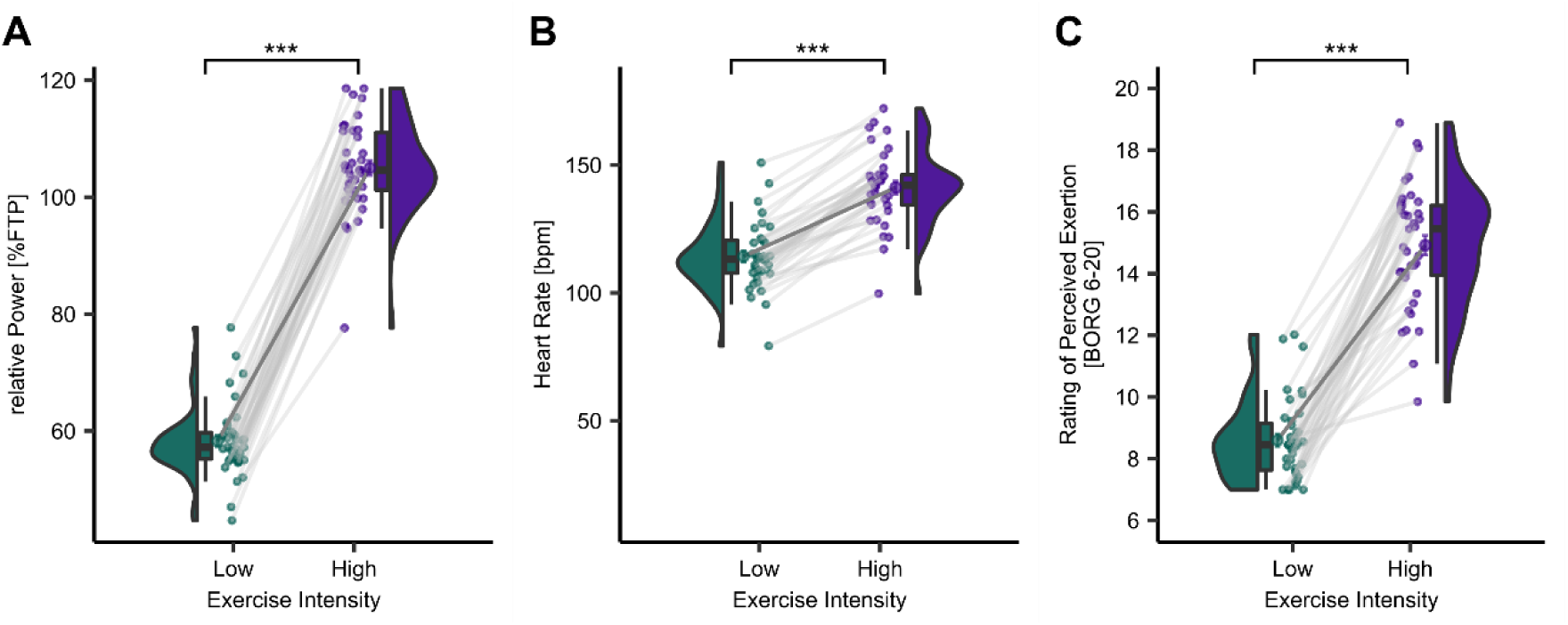
Successful implementation of high-(HI) and low-intensity (LI) exercise. (**A**) Relative Power (%FTP), (**B**) heart rate in beats per minute (bpm), and (**C**) rating of perceived exertion (RPE; BORG scale) during low-(LI; green) and high-intensity (HI; purple) cycling were all significantly different. *P*-values were calculated using a paired *t*-test (two-tailed, power: *N* = 38, heart rate: *N* = 34, BORG: *N* = 39). * *P* < 0.05, ** *P* < 0.01, *** *P* < 0.001.

### The effect of drug treatment in response to stimulus intensity

Next, we investigated the effect of our drug treatment with the µ-opioid antagonist naloxone (NLX). Previous studies have shown that NLX can increase perceived pain by blocking the effect of endogenous opioids (Eippert et al., 2009; Schoell et al., 2010). In line with those findings, we observed a significant interaction of drug and stimulus intensity on heat pain ratings (*β* = 0.10, CI [0.02, 0.18], *SE* = 0.04, *t*(2755) = 2.46, *P* = 0.01; Figure 4A, Table S5, and S6), where, with increasing stimulus intensities the differences between ratings in the NLX and SAL condition increased (*β* = 2.00, *CI* [0.40, 3.61], *SE* = 0.81, *t*(77) = 2.46, *P* = 0.02; Figure 4B, Table S7). We also investigated this interaction of drug and stimulus intensity between sexes and observed this interaction in females (*β* = 0.17, *CI* [0.06, 0.28], *SE* = 0.06, *t*(1471) = 3.00, *P* = 0.003; Figure 4C and Table S8) but not in males (*β* = 0.02, *CI* [−0.09, 0.13], *SE* = 0.06, *t*(1255) = 0.32, *P* = 0.75; Figure 4D and Table S9). The difference in pain ratings between the drug treatment conditions revealed a significant main effect of stimulus intensity (*β* = 3.41, *CI* [1.27, 5.54], *SE* = 1.09, *t*(76) = 3.12, *P* = 0.003) and sex (*β* = 9.75, *CI* [1.23, 18.23], *SE* = 4.42, *t*(101.60) = 2.21, *P* = 0.03; Figure 4E and Table S10). In contrast to females, males showed an overall large drug effect at every stimulus intensity. These findings demonstrate a robust drug effect, where pain ratings increased by blocking µ-opioid receptors. The increasing magnitude of the drug effect with increasing stimulus intensities indicates that the opioidergic system might be more extensively employed by more intense noxious stimuli. Since this magnitude differs for lower intensities between males and females it also indicates a possible sex-dependent effect regarding endogenous opioids.

**Figure 4.**
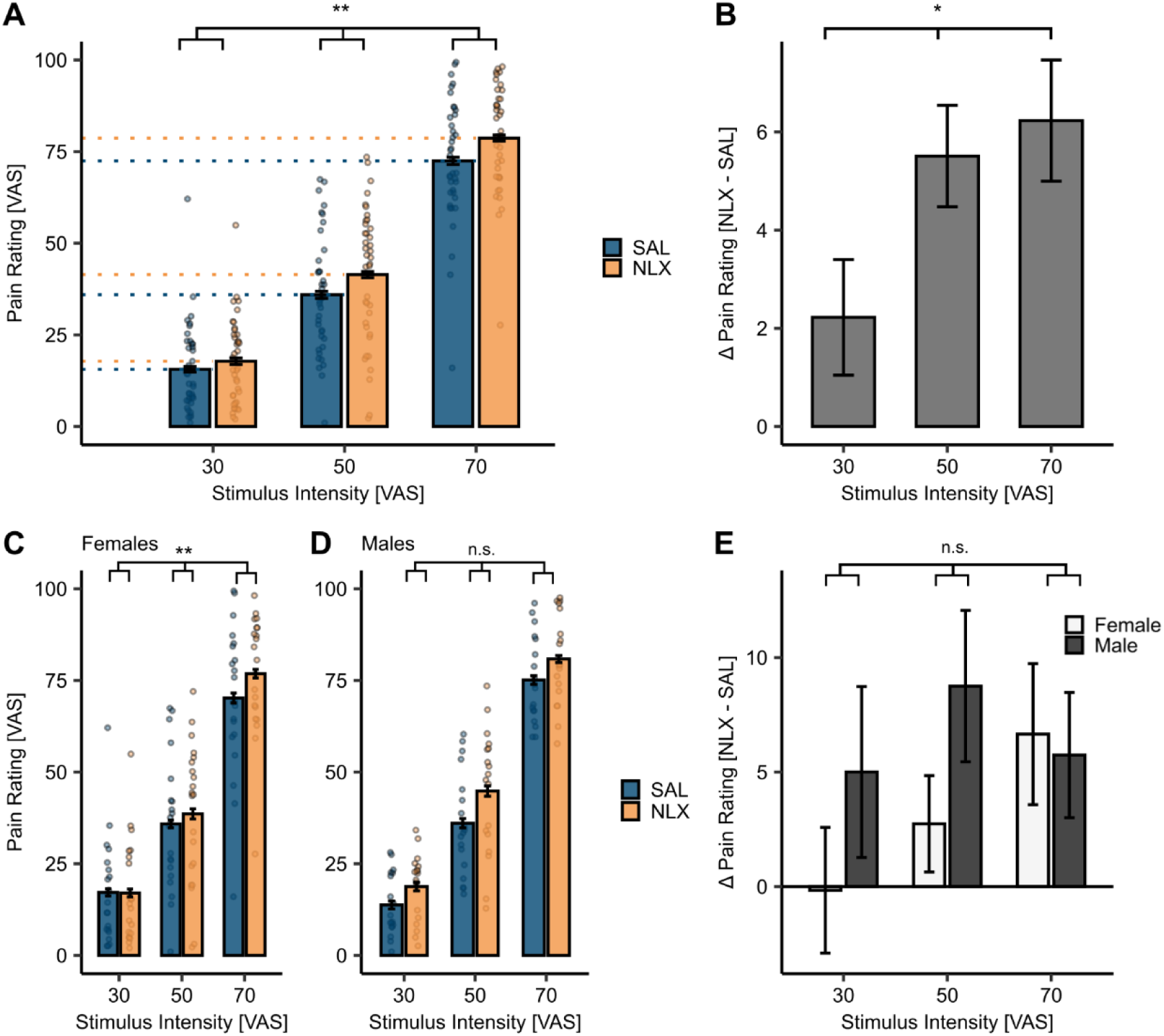
Behavioural results for the effect of drug treatment on pain. (**A**) Heat pain ratings revealed a significant interaction between drug and stimulus intensity (*P* = 0.01). Mean heat pain ratings were significantly higher under naloxone NLX treatment (orange) compared to placebo (SAL) (blue) across stimulus intensities. The *P*-value indicates a significant interaction effect of stimulus intensity and drug. (**B**) Differences in heat pain ratings between NLX and SAL condition (NLX – SAL) at each stimulus intensity revealed a significant main effect of stimulus intensity (*P* = 0.02). (**C-D**) Heat pain ratings at all stimulus intensities in both drug treatment conditions are significantly higher in NLX (orange) compared to SAL (blue) conditions for both sexes. In females (C), a significant interaction of stimulus intensity and drug was evident (*P* = 0.003) but not in (D) males (*P* = 0.75). (**E**) Differences in heat pain ratings between NLX and SAL condition (NLX – SAL) at each stimulus intensity for females (white) and males (dark grey) revealed a significant main effect of stimulus intensity (*P* = 0.003) and sex (*P* = 0.03) with an interaction showing a trend (*P* = 0.06). *P*-values were calculated using the LMER model for the interaction of stimulus intensity and drug. * *P* < 0.05, ** *P* < 0.01, *** *P* < 0.001. Error bars depict the SEM (*N* = 39).

In the next step, we identified brain regions associated with pain processing and opioidergic pain modulation. We observed a significant activation (contrast interaction of stimulus intensity and drug treatment) in the PAG (MNI*_xyz_*: −2, −24, −8, T = 5.17, *P_corr-SVC_* = 0.01; Figure 5A). The extracted parameter estimates from the peak voxel showed that NLX increased activation compared to SAL, especially for the highest stimulus intensity (Figure 5B and 5C). We used post-stimulus averaging (Finite Impulse Response or FIR model) to extract the mean time course for the highest-intensity stimulus (VAS 70). A significant difference in activation was evident seven seconds after stimulus onset (BOLD response) for the remainder of the stimulus in the peak voxel in the PAG (Figure 5D; for an uncorrected map at *P_uncorr_* < 0.001 see Figure S4). We also investigated the contrast heat NLX > heat SAL where activation in the right antIns showed a trend at whole-brain correction (MNI*_xyz_*: 46, 8, 6, T = 5.70, *P_corr-WB_* = 0.08; Figure S5).

**Figure 5.**
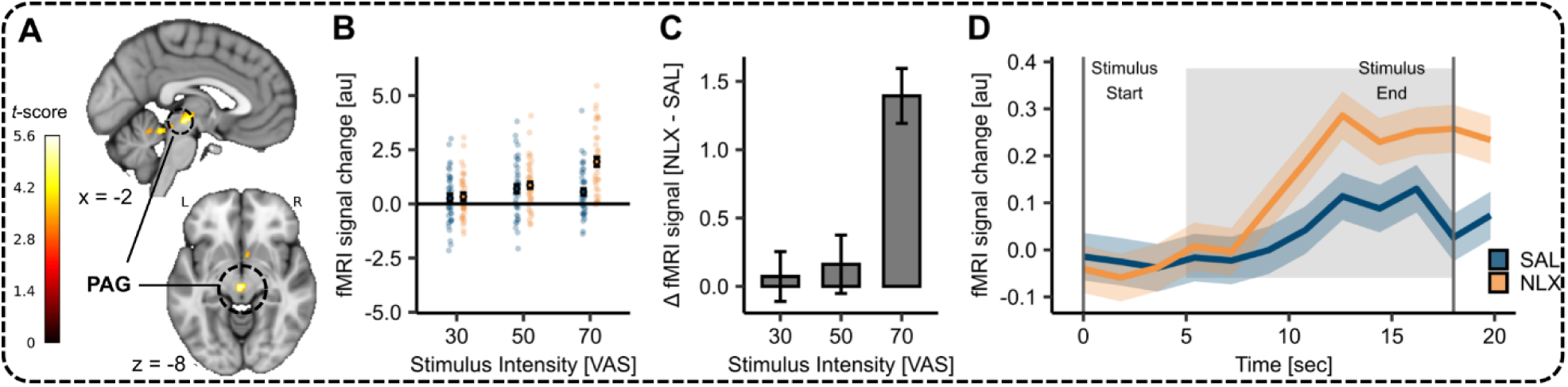
Effect of NLX in the periaqueductal grey (PAG). (**A**) Activation for the interaction of stimulus intensity and drug in the PAG (MNI_xyz_: −2, −24, −8, T = 5.17, *P_corr-SVC_* = 0.01) superimposed onto an MNI template brain. (**B**) Parameter estimates for SAL (blue) and NLX (orange) conditions for the respective peak voxel in the PAG (MNI_xyz_: −2, −24, −8). (**C**) Difference between parameter estimates of the peak voxel between NLX and SAL condition at each stimulus intensity. (**D**) Time course of BOLD responses for SAL (blue) and NLX (orange) during high pain (VAS 70) in this voxel. The shaded areas depict the SEM. The grey solid lines indicate the start and end of the painful stimulus. The shaded grey area displays the approximate time window for BOLD response taking into account a 5s delay of the hemodynamic response function.

### No difference between HI and LI aerobic exercise on pain modulation

After we established a successful exercise and drug intervention, we investigated the modulating effect of exercise intensity on pain ratings and neuronal responses in the SAL condition. There was no main effect of exercise intensity on pain as there was no hypoalgesic effect evident in the behavioural pain ratings comparing HI to LI exercise in the SAL condition (*β* = 1.19, *CI* [−1.85, 4.22], *SE* = 1.55, *t*(1354) = 0.77, *P* = 0.44; Figure 6A, blue bars and Table S11). This contrasts our preregistered hypothesis. Furthermore, there were no significant differences in BOLD activation between both exercise conditions (contrasts: exercise high > low (SAL) or exercise low > high (SAL)). For the uncorrected activation maps at *P_uncorr_* < 0.01 please refer to Figures S6 and S7. Additionally, we extracted the parameter estimates from the preregistered ROIs (RVM, PAG, and frontal midline; Figure 6B-D) and estimated the LMER models with exercise intensity on parameter estimates in the SAL condition (Figure 6E-G, blue bars). None of these models yielded a significant main effect of exercise intensity (Tables S12-S14).

**Figure 6.**
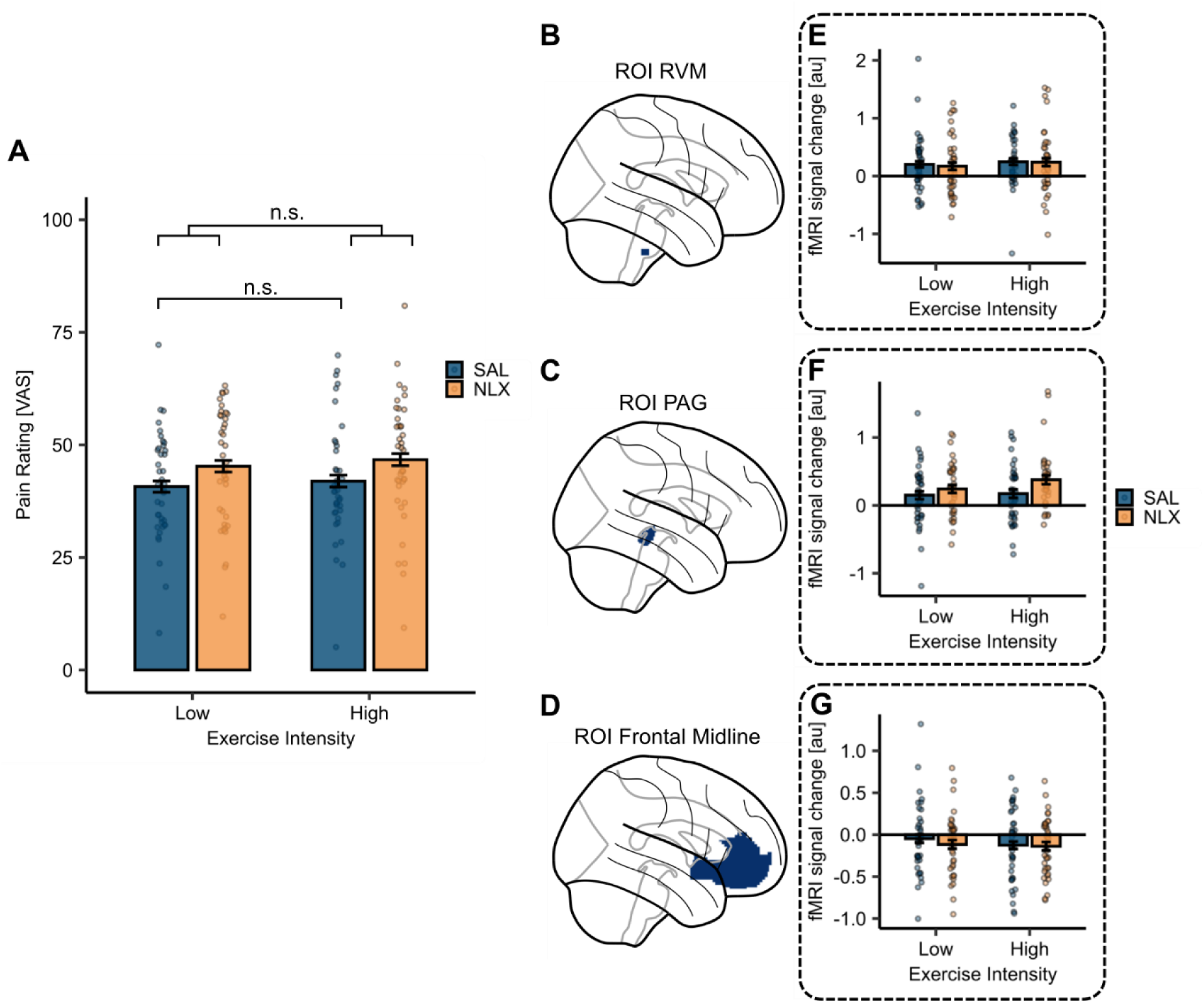
Effect of exercise intensity and drug treatment on pain modulation. (**A**) No significant main effect of exercise intensity on pain ratings in the SAL condition (*P* = 0.44, blue bars). The *P*-value was calculated using the LMER model with exercise intensity. In a separate LMER model for the interaction of exercise intensity and drug treatment, this interaction effect was not significant (*P* = 0.91) but a significant main effect of drug treatment (*P* = 0.005) was evident. Bars depict the average pain ratings in the SAL (blue) and NLX (orange) conditions in both exercise conditions averaged across all stimulus intensities and the dots represent the subject-specific average ratings averaged across all stimulus intensities. Error bars depict the SEM (*N* = 39). (**B**-**D**) Regions of interest (ROIs) in the (B) RVM, (C) PAG, and (D) frontal midline (comprised of ACC and vmPFC). (**E**-**G**) Parameter estimates extracted from both exercise and treatment conditions for the respective ROIs showed no significant effect of exercise intensity in the SAL condition (Tables S12 – S14) as well as no interaction of stimulus intensity with drug treatment (Tables S16 – S18). n.s. = not significant, * *P* < 0.05, ** *P* < 0.01, *** *P* < 0.001.

Despite not detecting a hypoalgesic effect of exercise intensity in the SAL condition, we investigated whether naloxone might alter the pain ratings depending on the exercise intensity. We estimated an LMER model with exercise intensity and drug treatment which yielded a significant main effect of drug treatment (*β* = 4.50, *CI* [1.36, 7.64], *SE* = 1.60, *t*(2755) = 2.81, *P* = 0.005) but no significant interaction effect of exercise intensity and drug treatment (*β* = 0.27, *CI* [−4.17, 4.71], *SE* = 2.27, *t*(2755) = 0.12, *P* = 0.91; Figure 6A and Table S15). For completeness, we have extended this model to include stimulus intensity, yielding no significant interaction of exercise intensity, drug treatment, and stimulus intensity (*β* = −0.05, *CI* [−0.20, 0.11], *SE* = 0.08, *t*(2751) = −0.56, *P* = 0.58; Figure S8). Corroborating this, there were no significant differences in BOLD activation for the contrasts interaction drug treatment and exercise intensity (positive and negative weights). For completeness, we visualised the uncorrected BOLD activation maps at *P_uncorr_* < 0.01 in Figures S9 and S10. We again extracted the parameter estimates from the preregistered ROIs (Figure 6B-D) across both treatment conditions and estimated the LMER models with the interaction of exercise intensity and drug treatment on parameter estimates from the respective ROIs (Figure 6E-G). None of these models yielded a significant interaction effect of stimulus intensity and drug treatment (Tables S16-S18). Since we could not establish a difference in the hypoalgesic effect of HI and LI aerobic exercise in the SAL condition, our preregistered hypothesis of reducing the hypoalgesic effect through naloxone administration behaviourally and neuronally could not be confirmed at either of the exercise intensities.

### Potential association of exercise-induced pain modulation with fitness level in the medial frontal cortex (mFC)

Although not preregistered, we additionally explored whether accounting for the different fitness levels of our participants in the statistical model can yield an exercise-induced analgesia effect. As a measure of fitness levels, we utilised the FTP (W/kg), and for ease of interpretation, we further used the difference scores between pain ratings after low-(LI) and high-intensity (HI) exercise bouts (LI – HI exercise pain ratings). Positive difference scores indicate hypoalgesia (i.e. higher pain ratings after LI exercise) whereas negative difference scores indicate hyperalgesia (i.e. higher pain ratings after HI exercise). This exploratory analysis showed a significant main effect of fitness level on differences in pain ratings in the SAL condition (*β* = 6.45, *CI* [1.25, 11.65], *SE* = 2.56, *t*(38) = 2.52, *P* = 0.02; Table S19) suggesting increased hypoalgesia with increasing fitness levels after HI compared to LI exercise, pooled across all stimulus intensities (Figure 7A). In the brain, the corresponding contrast (exercise high > exercise low with the mean-centered covariate FTP (W/kg)) was also investigated in the SAL condition. Here we observed a significant activation of the medial frontal cortex (mFC; MNI_xyz_: 6, 45, 10; T = 4.59, *P_corr-SVC_* = 0.05; Figure 7B). The parameter estimates from this voxel revealed that an increase in fitness level was negatively associated with the difference scores between parameter estimates from the drug treatment conditions (Figure 7C). These results suggest that the extent and direction of exercise-induced pain modulation potentially depend on the individual fitness levels and might be mediated by the mFC. (See Figure S11 for pain ratings and parameter estimates separately for HI and LI exercise and Figure S12 for an uncorrected activation map at *P_uncorr_* < 0.001).

**Figure 7.**
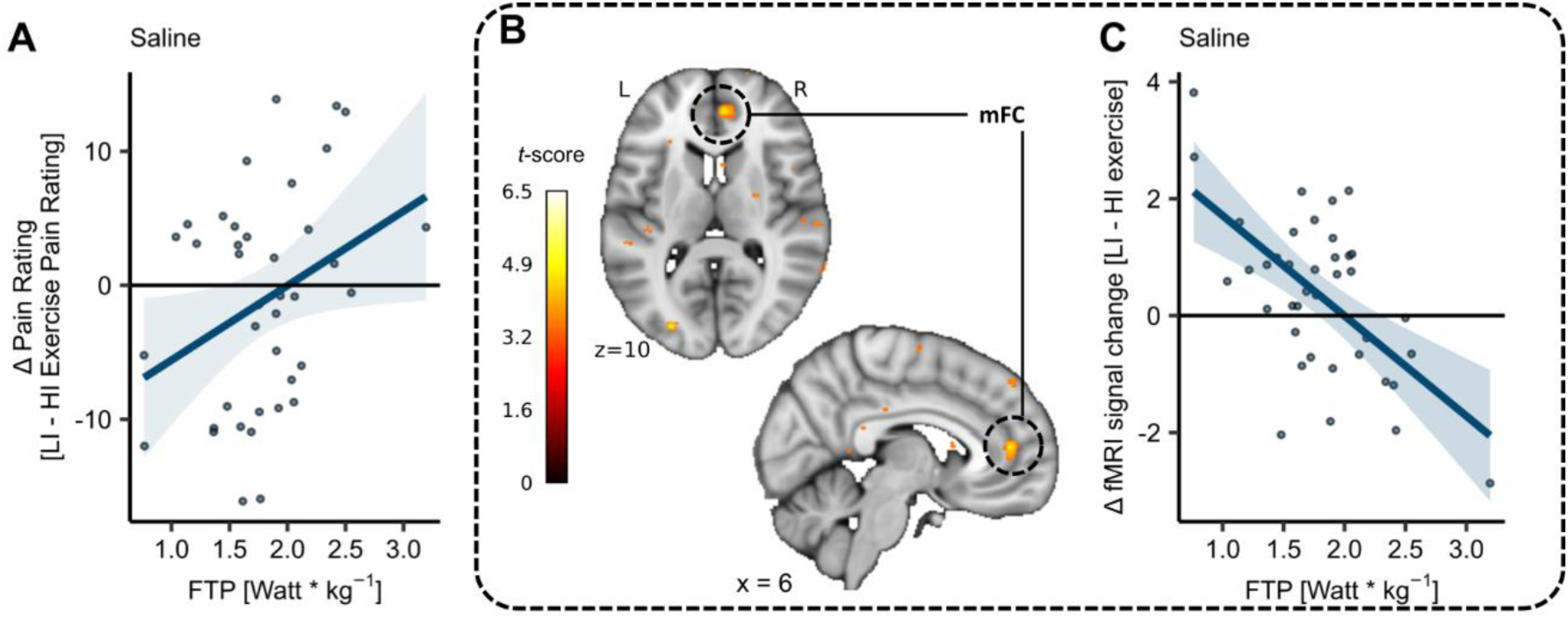
Fitness level on difference pain ratings (LI-HI exercise) and mFC activation. (**A**) Subject-specific differences in heat pain ratings (dots) between low-intensity (LI) and high-intensity (HI) exercise conditions (LI – HI exercise pain ratings) and corresponding regression line pooled across all stimulus intensities in the SAL condition. Fitness level (FTP) showed a significant positive relation to heat pain ratings with a significant main effect of FTP (*P* = 0.02) on difference ratings. (**B**) Cortical activation for contrast: exercise high > exercise low with mean-centred covariate FTP (weight-corrected) in the right mFC (MNI_xyz_: 6, 45, 10; T = 4.59, *P_corr-SVC_* = 0.05) across all stimulus intensities in the SAL condition superimposed onto an MNI template brain. (**C**) Differences between parameter estimates of LI and HI exercise conditions (LI – HI exercise parameter estimates) from respective peak voxel, plotted for each subject as a function of fitness level (FTP). Regression lines are visualised and shaded areas represent the SEM (*N* = 39).

### Exploring the effect of sex, fitness level, and endogenous opioids on exercise-induced pain modulation along the mFC

As a second, not preregistered, exploratory analysis we tested for sex effects in exercise-induced pain modulation. This was motivated by our preceding analyses suggesting that males and females differ in the modulation of pain by NLX administration at different stimulus intensities (Figure 4E). We observed a significant interaction of sex, fitness level, and drug (*β* = −13.12, *CI* [−23.69, −2.56], *SE* = 5.42, *t*(190) = −2.43, *P* = 0.016, Figure 8A and Figure 8B; Table S20). In the SAL condition, males showed larger hypoalgesia with increasing fitness levels (Figure 8A, red line) whereas females showed no hypoalgesic response regardless of fitness levels (Figure 8A, blue line). In the NLX condition, however, there was no significant association between fitness levels and difference in pain score in males (Figure 8B, red line) but a small positive correlation in females (Figure 8B, blue line). These results possibly hint at diverging pain modulatory mechanisms in males and females depending on their fitness level and drug treatment. In the next step, we investigated this interaction of fitness level, sex, and drug in the neuroimaging data, formally implemented as a two-sample (male vs. female) *t*-test for the interaction contrast between exercise intensity and drug with the subject covariate of individual FTP. This revealed significant activation in the mFC (MNI_xyz_: 12, 64, 2; T = 4.78, *P_corr-SVC_* = 0.039; Figure 8C). The parameter estimates from the peak voxel are shown for both drug treatment conditions and both sexes in Figures 8D and 8E. In the SAL condition, males showed a negative association of the difference (EIH) parameter estimates with fitness level (Figure 8D, red line), whereas females showed a positive association (Figure 8D, blue line). Importantly, this pattern was abolished under NLX (Figure 8E; see Figure S13 for an uncorrected activation map at *P_uncorr_* < 0.001). Despite being an exploratory analysis, these findings suggest an interaction between sex and fitness level in opioidergic mechanisms that might mediate exercise-induced pain modulation.

**Figure 8.**
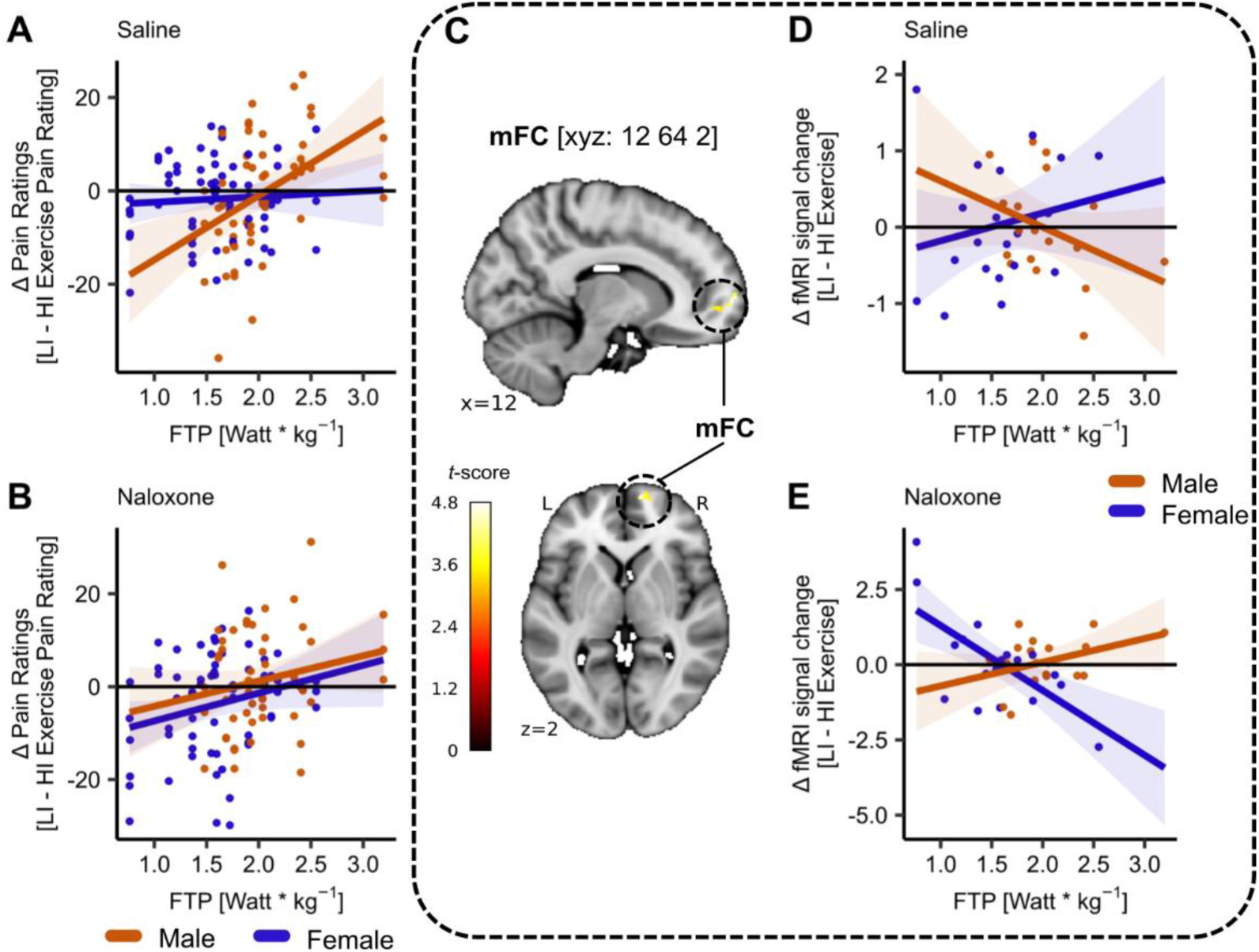
Exercise-induced pain modulation potentially depends on sex, fitness level, and drug treatment in mFC. (**A**) Exercise-induced pain modulation in the SAL condition for males (red) and females (blue). Males showed larger hypoalgesic responses with increasing fitness levels as indicated by positive difference ratings in the SAL condition. Females showed no association between fitness levels and difference ratings. (**B**) No exercise-induced pain modulation after blocking µ-opioid receptors with NLX in males (red) and females (blue). (**C**) The activation pattern in the mFC (MNI_xyz_: 12, 64, 2; T = 4.78, *P_corr-SVC_* = 0.039) resulting from a two-sample *t*-test (two-tailed) between males and females for contrast interaction of exercise and drug with covariate FTP superimposed onto an MNI template brain. (**D-E**) The difference in parameter estimates from the peak voxel in the mFC in (D) SAL and (E) NLX condition for males (red) and females (blue). Each dot represents the difference in parameter estimates between LI and HI exercise conditions (LI – HI exercise) for each subject averaged across all stimulus intensities. Shaded areas represent the SEM (female: *N* = 21, male: *N* = 18).

## Discussion

This double-blind cross-over study aimed to disentangle the neural and pharmacological mechanisms of exercise-induced pain modulation in healthy individuals of varying fitness levels. There was no main effect of exercise-induced pain modulation after high-intensity (HI) exercise compared to low-intensity (LI) exercise in the saline (SAL) condition, conflicting our preregistered hypothesis. What is more, naloxone (NLX) administration resulted in increased pain ratings across both exercise conditions but showed no interaction with exercise intensity. These results suggest that there is no difference between HI and LI aerobic exercise on heat pain and, further, no direct involvement of the endogenous opioid system. In an exploratory analysis, we observed a significant interaction involving fitness level, sex, and drug administration where males exhibited greater hypoalgesia, particularly as fitness levels increased under the saline (SAL) condition. This effect was diminished upon administration of naloxone (NLX) and was also apparent in opioidergic regions of the pain modulatory system, namely the medial frontal cortex (mFC).

As expected, we were able to successfully induce heat pain at different stimulus intensities, mirrored by different levels of activation in the right anterior insula (antIns), right dorsal posterior insula (dpIns), and right midcingulate cortex (MCC). As part of a highly distributed pain processing system (Coghill, 2020; Tracey and Mantyh, 2007) the antIns has been established to encode pain intensity (Coghill, 2020) and salience (Wiech et al., 2010) whereas the dpIns has been shown to encode stimulus intensity and salience specific to heat pain (Horing et al., 2019; Segerdahl et al., 2015). The function of the MCC has been more equivocal since some studies attributed a domain-general function of integrating pain, negative affect, and cognitive control (Shackman et al., 2011) whereas others argued in favour of a domain-specific representation of painful stimulation (Kragel et al., 2018).

### Opioidergic mechanisms in heat pain

Previous research has shown that heat pain is modulated by the descending pain modulatory system including the PAG (Borras et al., 2004; Petrovic et al., 2002) mainly through opioidergic mechanisms such as placebo analgesia (Eippert et al., 2009; Petrovic et al., 2002; Tinnermann et al., 2022). Accordingly, studies showed VAS ratings of intense heat stimuli to be increased after NLX administration (Borras et al., 2004; Kut et al., 2011), along with a significant modulation of BOLD responses in the PAG (Eippert et al., 2009). In line with this, we could show that NLX significantly increased perceived painfulness mirrored by activation in the PAG (Corder et al., 2018). An important aspect of our study is that the magnitude of opioidergic pain modulation increased with pain intensity in females, whereas males showed a constant, but high magnitude of pain modulation across stimulus intensities when NLX was administered. This finding suggests two things: firstly, endogenous pain modulation through µ-opioid receptors increases with increasing pain intensity and, secondly, the magnitude of this modulation differs between sexes. Since our study employed a within-subject cross-over design with an established pharmacological intervention (Leknes and Atlas, 2020; Trøstheim et al., 2022), we ensured our results were unbiased and highly robust.

### No difference between HI and LI aerobic exercise on pain modulation

We did not detect significant differences in pain ratings and BOLD signal between the exercise conditions (HI and LI) in the SAL condition. Several factors might contribute to this null finding. For one, the exercise intensities employed in this study might not have been suitable to induce pain modulation. Most studies using peak oxygen consumption (VO_2max)_ (Koltyn et al., 1996; Micalos and Arendt-Nielsen, 2016; Vaegter et al., 2014) or the maximum heart rate (HR_max_) (Gomolka et al., 2019; Gurevich et al., 1994; Jones et al., 2017; M. D. Jones et al., 2019; Vaegter et al., 2014) as a measure of exercise intensity were able to induce hypoalgesia with HI exercise ranging between 65%-75% VO_2max_ and above 70% HR_max_. Previous research has further suggested that HI exercise produces greater hypoalgesia compared to LI exercise (60-70% HR_max_ vs. light activity: M. D. Jones et al., 2019; 70% vs. 50% HR_max_: Naugle et al., 2014; 75% vs. 50% VO_2max_: Vaegter et al., 2014). What is more, previous studies have often employed a resting control condition, which somehow limits the comparability of exercise conditions since a resting baseline does not control for unspecific factors such as cognitive pain modulation (i.e. attentional load or distraction) that is mediated by endogenous opioids (Brooks et al., 2017; Sprenger et al., 2012) and, thus, potentially exercise. Furthermore, different aerobic exercise durations have been shown to induce hypoalgesia with durations ranging between 8 minutes to 2 (for review see Koltyn (2002)). Previous studies were able to induce hypoalgesia at 10-15 minutes of HI aerobic exercise (75% VO_2max_: Gomolka et al., 2019; 63% VO_2max_: Gurevich et al., 1994; self-paced: Haier et al., 1981; 60-70% HR_max_: M. D. Jones et al., 2019; 85% HR_max_: Sternberg et al., 2001; 75% VO_2max_: Vaegter et al., 2015). Thus, we opted for the control condition of LI exercise instead of rest to also add to the knowledge of a dose-response relationship of exercise intensity needed to induce a hypoalgesic effect. We confirmed the intensities of both exercise conditions by physiological (heart rate (HR)) and psychophysiological measures (rating of perceived exertion (RPE)). Participants indicated significant differences in HR and RPE between both exercise conditions, confirming the actual and perceived distinction between the exercise conditions.

Another factor to consider is that our study utilised the FTP_20_ test as a tool to determine the highest power output a cyclist can maintain in a quasi-steady state (Allen and Coggan, 2006). It is frequently used in studies investigating athletes (Borszcz et al., 2020; Mackey and Horner, 2021; Mc Grath et al., 2019) and shows a high correlation with other measures such as the FTP_60_ test (*r* = 0.88) (Borszcz et al., 2018). Compellingly, the FTP_20_ test does not rely on approximations or self-reports but on the actual power maintained (Allen and Coggan, 2006). Previous reviews have shown that hypoalgesia following exercise is most reliably induced by HI (i.e., > 75% VO_2max_) exercise (Koltyn, 2002; Naugle et al., 2012; Wewege and Jones, 2020). Thus, we decided to employ a HI exercise protocol using the FTP threshold as a reliable measure. Nevertheless, since our study is one of the first studies to employ this FTP_20_ test with a non-athlete, balanced sample of varying fitness levels, one could argue that the FTP_20_ test is not a reliable estimation of the HI exercise condition. Despite research showing that the FTP is a reliable marker of VO_2max_ (Denham et al., 2020), we did not directly measure blood lactate levels and VO_2max_ during the exercise and, thus, did not control for the intensity domains (i.e., moderate – heavy – severe). Since the FTP reflects the highest possible intensity by which a steady-state VO2 is maintained (Sørensen et al., 2019), the FTP might be a demarcation point between the heavy and severe domains (A. M. Jones et al., 2019); although this has been criticised (Wong et al., 2022). Consequently, some participants might have exercised in the severe domain during the HI exercise and in the moderate domain during the LI exercise. However, as mentioned above, the RPE and relative power (%FTP) still confirmed a successful implementation of the exercise intervention at the anticipated intensities suggesting the FTP_20_ test to be a reliable estimation of functional threshold power in our sample. Overall, we ensured the highest accuracy in determining the individual exercise intensities at HI and LI.

Furthermore, the composition of the sample, which included non-athletes with varying fitness levels and of both sexes, might have contributed to the missing difference between HI and LI aerobic exercise on pain modulation. In human studies, many findings on exercise-induced pain modulation are derived from studies in athletes or physically active individuals showing increased hypoalgesia following exercise compared to subjects with lower fitness levels (Crombie et al., 2018; Geva and Defrin, 2013; Naugle and Riley, 2014; Schmitt et al., 2020). This observation might be related to greater exercise-induced opioid release in structures such as the PAG or the frontal medial cortex (including the ACC) in fitter individuals (Saanijoki et al., 2022; Sluka et al., 2018). Furthermore, exercise-induced analgesia in rodents and humans has predominantly been studied in all-male samples (Koltyn et al., 2014; Mogil, 2012; Niesters et al., 2010). Of those studies that have considered potential sex differences in exercise-induced pain modulation, some (Brellenthin et al., 2017; Koltyn et al., 2013; Naugle et al., 2014) could not reveal a sex effect, whereas others (Koltyn, 2000; Sternberg et al., 2001; Vaegter et al., 2014) revealed a stronger effect in females. This is in contrast to results in other domains, where opioid-dependent pain relief through conditioned pain modulation (Bulls et al., 2015; Ge et al., 2004; Granot et al., 2008) or placebo analgesia (Aslaksen et al., 2011; Desai et al., 2022) has been observed to be stronger in males. We deliberately did not employ professional athletes in this study but rather individuals of varying fitness levels of both sexes to make our results more applicable to the general population.

In the context of exercise-induced pain modulation, different types of pain have been investigated (i.e. thermal, pressure, and electrical) as well as different measurements of such (threshold, intensity, and tolerance) (for reviews see Koltyn, 2000; Naugle et al., 2012; Vaegter & Jones, 2020). For aerobic exercise, the highest effect sizes were reported for pressure pain (pain intensity: Cohen’s *d* = 0.69) closely followed by heat pain (pain intensity: *d* = 0.59) (Naugle et al., 2012). Despite having successfully induced pressure pain using computer-controlled cuff pressure algometry, we were unable to detect significant changes in pain ratings for different exercise intensities. However, most of the studies employing pressure pain used handheld algometry exerting local pressure which can be prone to experimenter bias (Vaegter and Jones, 2020) and those results obtained might not apply to cuff pressure pain. Thermal heat pain was the other chosen method to induce pain and, as mentioned above, we were able to successfully induce heat pain at different stimulus intensities, mirrored by different levels of activation in the right antIns, right dpIns, and right MCC. Since our method to induce thermal heat pain corresponds to the method used in previous studies, the results of this pain modality might be more comparable to those of existing studies.

### Exploring the effect of fitness level and sex on exercise-induced pain modulation

In the exploratory analyses we identified that pain decreases following HI compared to LI exercise to be associated with higher individual fitness levels in the SAL condition. In agreement with previous studies (Geva and Defrin, 2013; Saanijoki et al., 2022; Schmitt et al., 2020; Tesarz et al., 2012), this suggests a dependence of exercise-induced pain modulation on individual fitness. Recent studies indirectly support this notion and observed higher self-reported physical activity levels to be associated with reduced temporal summation in heat pain (Naugle and Riley, 2014) and increased heat pain thresholds to be associated with individual fitness levels in males (Schmitt et al., 2020). In a larger cohort, an association between physical activity levels and pressure pain tolerance (Årnes et al., 2023) as well as cold pressor pain tolerance (Årnes et al., 2021) was observed. Along with our behavioural findings, we identified increased activation in the mFC after HI exercise to be associated with higher fitness levels in the SAL condition. Previous research has reported increased activation in the rACC to be associated with opioid analgesia (Petrovic et al., 2002; Tinnermann et al., 2022), thus implying opioidergic involvement of this structure in mediating exercise-induced pain modulation (Petrovic et al., 2002). Adding to this, the pregenual ACC showed decreased activation in response to painful heat following a 2-h run compared to before the run in trained male elite runners (Scheef et al., 2012). Furthermore, larger decreases in µ-opioid receptor binding in regions including the ACC, bilateral insula, and OFC in healthy males have been observed to be associated with higher self-reported physical fitness levels (Saanijoki et al., 2022). This finding could support observations showing that repeated exercise “trains” the mFC, which has a high µ-opioid receptor density (Corder et al., 2018) and thus enables exercise-induced pain modulation even in the context of brief bouts of HI exercise.

Furthermore, we explored the interaction between sex, fitness level, and drug, where males exhibited larger hypoalgesia following HI exercise with increasing fitness levels, which was attenuated by NLX. This pattern was not evident in females. Our brain data also showed this three-way interaction in the mFC. More precisely, in the SAL condition, males showed higher activation after HI exercise compared to LI exercise in association with higher fitness levels, and this pattern was reversed in the NLX condition. Females showed the reversed pattern to males in both, SAL and NLX, conditions. Since rodent and early human research have also predominantly employed male samples to avoid hormonal-related fluctuations (Mogil, 2012; Mogil and Chanda, 2005), this might have led to an over-generalisation of opioidergic effects (Koltyn et al., 2014; Mogil, 2012) whilst neglecting sex differences in opioid analgesia (Bodnar and Kest, 2010; Niesters et al., 2010). Hence, it comes as no surprise that findings in humans are heterogeneous (Koltyn, 2000; Wewege and Jones, 2020) with some studies showing no difference in exercise-induced pain modulation between males and females (Koltyn et al., 2014, 2013; Smith, 2004) whereas other studies showed hypoalgesia from exercise to be greater in women (Vaegter et al., 2014) but most research on exercise-induced pain modulation has been conducted in men. The few existing imaging studies show that frontal midline structures (ACC, OFC) with high opioid receptor concentration are involved in exercise-induced (pain) modulation (Boecker et al., 2008; Saanijoki et al., 2022; Scheef et al., 2012). A recent meta-analysis has shown that increases in the medial frontal cortex (including the vmPFC) activity along with reduced activation in regions associated with noxious stimuli (i.e. ACC) could be linked to an expectation of reduced pain, in this case, after HI exercise (Atlas and Wager, 2014). When comparing athletes and non-athletes, one study identified an interaction of sex and athletic status where female athletes displayed significantly lower cold pressor ratings than female non-athletes, but there was no difference in men (Smith, 2004). These exploratory findings suggest sex-dependent and fitness level-dependent effects in exercise-induced pain modulation but do not provide a definite answer. Future research should further investigate the role of sex and fitness levels concerning exercise-induced hypoalgesia, specifically investigating females with higher fitness levels.

### Limitations

One limitation that should be considered when interpreting the results is the influence of other mediating factors in exercise-induced pain modulation such as endocannabinoids (eCB; Crombie et al., 2018; Siebers et al., 2021). Previous animal studies have established their crucial influence in modulating mood and promoting anxiolysis following exercise (Fuss et al., 2015). In humans, however, the interpretability of results remains limited since only peripheral eCB can be measured and central eCB blockage is difficult.

In conclusion, our study showed no difference in pain modulatory effects between HI and LI aerobic exercise, neither behaviourally nor in brain responses. However, explorative analyses suggested that there is, in fact, an interaction between sex and fitness level in mediating exercise-induced pain modulation through opioidergic mechanisms in the mFC. To thoroughly investigate this, future research should specifically address the link between endogenous opioids, fitness levels, and sex-dependent differences in exercise-induced pain modulation.

## Materials and Methods

The study was preregistered with the WHO-accredited *Deutsches Register für Klinische Studien* (DRKS; Study ID: DRKS00029064; German Registry for clinical studies) before data acquisition.

### Experimental Design

This randomised control fMRI study employed a pharmacological intervention with the µ-opioid antagonist NLX in a within-subject design and was divided into a calibration and two experimental days (Figure 1). On the experimental days, participants cycled outside the MR for 10 minutes at HI or LI on a stationary bike per block. Afterward, they rated the perceived painfulness of heat and pressure stimuli inside the MR. This procedure was repeated for four blocks per experimental day. The objective of this study was to identify the behavioural, pharmacological, and neuronal underpinnings of exercise-induced pain modulation.

### Participants

Forty-eight healthy, right-handed participants were invited to take part in the study. In the framework of the within-subject design, all participants underwent all conditions which included an exercise factor (high vs. low exercise intensity) and a pharmacological intervention (NLX vs. SAL). All participants received the pharmacological intervention in a randomised double-blind fashion on separate study days, where the µ-opioid antagonist NLX or placebo SAL was administered i.v. Our final sample size was determined based on previous behavioural studies investigating exercise-induced pain modulation after aerobic exercise in a pre-post measurement design across different pain modalities (*d* = 0.59 for heat pain intensity, *d* = 0.69 for pressure pain intensity) (Naugle et al., 2012). According to this, a sample size of *N* = 39 was sufficient to reveal results at effect size *f* = 0.3, (via G*Power 3.1, *α* = .05, 1-*α* = 0.95, 1 group, 2 measurements with correlation *r* = 0.5). Based on previous fMRI studies it can also be assumed that a sample size of *N* = 30 would be sufficient to detect changes in the brain (Mumford, 2012). In this study, the term “sex” refers to “sex assigned at birth” as indicated by self-reports from the participants. Multiple options (“male”, “female”, “other”, and free text field) were provided. All participants reported having been assigned the “female” of “male” sex at birth.

### Exclusion of participants

Upon entering the study, participants were screened for anxiety (State and Trait questionnaires of the German short version of the State-Trait Anxiety Inventory; STAI) (Laux et al., 1981), depression (Beck Depression Inventory 2; BDI-II) (Beck et al., 2011) as well as potential MR contraindications. Furthermore, participants’ body mass index (BMI) was required to range between 18 and 30. On the first day of the study, an electrocardiogram (ECG; cardiofax, Electrocardiograph-1250, Nihon Koden) was recorded to prove no cardiovascular irregularities. The age of participants was between 18 and 45 years. Eight participants were excluded from the study for the following reasons. Two participants withdrew from the study after the first day due to circumstantial reasons (moving, no appointment found). Two participants were excluded after day one due to an unreliable thermal calibration and a BMI score below 18, respectively. Two participants did not wish to continue the study after day two due to personal reasons. Two participants were excluded after day two due to a syncope when administering the drug (in both cases SAL has been administered). Upon data screening, one subject was excluded due to excessive movement (> 0.6 mm difference between volumes within runs in more than 5% of volumes in all runs) in the MR scanner. A total of thirty-nine participants (Age: *M* = 26.03, *SD* = 4.8, 21 female) were included in the final sample (Table S21).

### Ethics approval

The study was approved by the Ethics committee of the medical board in Hamburg (’Aerztekammer’) and conducted following the Declaration of Helsinki (World Medical Association, 2013). All subjects provided informed consent upon entering the study after having been informed about the study procedures including thermal and pressure stimulation, the MR procedure, the double-blind nature of the pharmacological intervention, and potential adverse effects of NLX by the study investigator and study physician. Upon completing or leaving the study, participants were informed of the order of pharmacological treatment received by the unblinded study physician.

### Thermal and Pressure Stimuli

Thermal stimulation was applied using a thermode (TSA-2, Medoc, Israel) attached to the left lower arm where four 2.5×2.5 cm squares were drawn below each other and numbered (Figure 1A). For the calibration, the thermode head was positioned in the second square. Each thermal stimulus lasted up to 17 seconds in total with a plateau of 15 seconds and a ramp speed of 13°C/s (ramp-up, ramp-down). For the pressure calibration, a computer-controlled cuff pressure algometer (CPAR; NociTech, Denmark, and Aalborg University, Denmark) consisting of a compressor tube and 13 cm wide tourniquet cuff (VBM medical, Sulz, Germany, 61 cm length) was used. The cuff was mounted to the left upper arm with a 3 cm distance to the cubital fossa to exert pressure on the upper arm tissue (Graven-Nielsen et al., 2015). Additionally, a protective tubular elastic dressing (Tricofix, D/5, 6 cmx20 cm) was positioned underneath the cuff to protect the bare skin from potential harm. The pressure was applied by inflating the chambers of the cuff with a maximal pressure limit of 100 kilopascal (kPa) and a rise speed dependent on the target pressure. The stimulus length of the pressure stimuli was also up to 17 seconds (ramp-up, 15s plateau, ramp-down).

### Calibration Procedure (Day 1)

The first day of the study took place outside the MR scanner and served as a calibration day (Figure 1A). Blood pressure, baseline heart rate (HR), and oxygen saturation (SPO2) were measured using a blood pressure cuff (boso medicus uno, bosch+sohn) and a pulse oximeter (pulox® Pusloximeter, Novidion GmbH), respectively. The thermal and pressure stimuli were calibrated individually but with an identical protocol in a pseudo-randomised order across participants. During calibration, participants remained in a supine position with the blinds down to emulate the inside of the MR scanner bore. As an initial step, participants received two low stimuli (10/20 kPa or 41°C/42°C) to get acquainted with the stimuli. Following this, the pain threshold was estimated by applying six adaptive stimuli which were rated on a binary scale (painful vs. not painful). The resulting pain threshold was used as a basis for the following linear regression algorithm aiming to collect ratings across different intensities on a Visual Analog Scale (10, 30, 50, 70, 90 VAS). After each stimulus, participants had to rate its painfulness (‘How painful was the last stimulus?”) on a VAS ranging from 0 (minimally painful) to 100 (almost unbearably painful) using the left and right key of a pointer as a button box (Logitech) with their right hand. The calibration was finished once sufficient rating coverage for all target intensities was reached. The thermal and pressure calibration aimed to identify super threshold intensities (°C and kPa) corresponding to 30, 50, and 70 on the VAS scale using this linear regression algorithm yielding comparable intensities across subjects.

### Functional Threshold Power Test (Day 1)

As a final step of the calibration day, a functional threshold power (FTP) test was conducted on a stationary cycle ergometer (KICKR Bike, Wahoo, Atlanta, United States). The FTP test was first described by Allen and Coggan (2006) and determines the maximum average power output that can be maintained for one hour which serves as an approximation of the anaerobic threshold (Allen and Coggan, 2012). The FTP_20_ test is derived from the original FTP test but estimates 95% of FTP based on 20 minutes of steady cycling at maximum possible power output. The test is divided into six stages lasting one hour in total (Figure 1A and Table S1). Within this test protocol, the average power output measured for the last 20-minute interval is taken as 95% of the actual FTP (Allen and Coggan, 2012). The FTP_20_ Test allows the identification of individually calibrated power zones. In this present study, we used HI exercise (Zone 4: Threshold, 91 - 106% of FTP) and LI exercise (Zone 1: Active recovery, ∼55% of FTP) as intensities. During the FTP_20_ test, participants were free to adjust the resistance of the ergometer whilst maintaining a constant cadence. Power output in Watts, heart rate (HR belt; Garmin® HRM-PRO™ PLUS) in beats per minute (bpm), and cadence (in RPM) were monitored throughout the FTP_20_ test. The current power and HR as well as the remaining time of each section were displayed on a 50-by-30 cm screen in front of the participants to allow for self-monitoring. Participants were asked to refrain from any exhausting physical activities the day of and before the calibration day. The FTP value from the FTP_20_ test was corrected for participants’ weight (FTP/weight) and served as an indicator of individual fitness level in the analyses. The distribution of the weight-corrected FTP for males and females is visualised in Figure S14. There was no significant association between heat pain thresholds and weight-corrected FTP (*r* = −0.23, *P* = 0.16; Figure S15). On the bike, the knee angle was adjusted to a 5-degree bend when the pedal stroke was at the bottom (Vaegter et al., 2019). The knee was also set to be over the pedal axle when the cranks were parallel to the ground. The upper body was adjusted to a forward lean with a slight bend of the elbows to maintain a neutral spine position with a natural lumbar flexion.

### Experimental Paradigm (Day 2 and Day 3)

The two experimental days were at least three days apart (*Median* = 7 days, *Mean* = 12.35 days, *SD* = 14.5 days) and identical with the only difference being the pharmacological treatment (NLX vs. SAL) received (Figure 1B). Each visit took place in the morning with a starting time of 8:15 a.m. and varied by a maximum of four hours between participants to account for potential circadian influences. Blood pressure and SPO_2_ were measured before each cycling block and monitored throughout the investigation. On each experimental day, a urine sample was tested for amphetamines, opiates, marijuana, methamphetamines, morphine (Surestep, Multidrug Test, Innovacon Inc., San Diego, USA), and potential pregnancies (only female participants; hCG Ultra Test, Serum/Urine 10mIU, Mexacare GmbH, Germany). Each experimental day consisted of 4 blocks. Each block included a 10-minute cycling block at a HI (100% FTP) or LI (55% FTP) in a pseudo-randomised order across participants (i.e., participants completed 2 cycling blocks at HI and 2 at LI exercise). The order of cycling blocks was kept constant across experimental days (Day 2 and Day 3). Each cycling block was immediately followed by an MR scan including the pain stimulation and ratings that lasted approximately 20 minutes. After this, participants’ HR and SPO_2_ were measured outside the MR (approximately 5-10 minutes) before the next block commenced starting again with 10-minute cycling. On the second experimental day (Day 3), an anatomical scan (T1) was acquired alongside conducting a re-calibration of thermal and pressure stimuli. The re-calibration was based on the previously calibrated intensities from the calibration day (Day 1) and was adjusted based on the ratings provided in the scanner corresponding to 30, 50, and 70 VAS. This procedure served to account for potential differences in perception due to the different environment of the MR scanner but also to acquaint participants with the procedure inside the MR to minimise the time of transition between the cycle ergometer and MR scanner.

### Drug administration

The procedure of drug administration and dosage was based on a previous study using NLX (Eippert et al., 2009) as well as a web-based application for detailed planning of µ-opioid antagonist administration (Trøstheim et al., 2022). Participants were positioned in a supine position before administering a bolus dose of NLX (0.15 mg/kg; Naloxon-ratiopharm® 0.4 mg/ml injection solution, Ratiopharm, Ulm, Germany) or SAL (Isotone Kochsalz-Lösung 0.9% Braun) via peripheral venous access (PVA) in the right antecubital vein. To ensure correct administration, two participants received intravenous access on the back of their right hand upon giving verbal consent and ensuring no pain was caused by the location of the intravenous line. Shortly after administering the bolus dose, the intravenous infusion dose of NLX (0.2 mg/kg/h diluted in SAL) or SAL was started using an infusion pump (Perfusor® Space, Braun, Munich, Germany). After ensuring that no pain was caused by the intravenous line and that participants felt comfortable, the first cycling block commenced. The first 10-minute cycling block also served to ensure that NLX reached a peak level in the blood plasma (Trøstheim et al., 2022) before positioning participants in the MR scanner. The infusion dose was constantly running throughout the whole duration of the study, including cycling blocks outside as well as pain stimulation inside the MR, providing a constant supply of NLX or SAL. Considering the relatively short half-life of NLX (∼70 minutes in blood plasma; Summary of Product Characteristics, Ratiopharm), this procedure ensures a steady-state concentration of NLX for the duration of the study (Trøstheim et al., 2022) with an effective central opioid block throughout the study (Eippert et al., 2009; Trøstheim et al., 2022). The experimenter (J.N.) and research assistants who interacted with the participants were blinded to the pharmacological intervention and remained blinded throughout the study. The research assistant administering the drug treatment was also blind to the treatment. Only the study physician (T. F.), who did not interact with the participants on the experimental days of the study was unblinded as to what drug was administered. Unblinding of the participants took place after the experiment by the study physician without anyone else present to prevent expectation induction in the experimenter and student assistants. Before unblinding the participants, they were asked what treatment they received on which experimental day (incorrect: *n* = 4, unsure/identical: *n* = 19, correct: *n* = 16).

### Questionnaires

On the calibration day (Day 1) participants completed the short form of the Profile of Mood States (POMS) (Curran et al., 1995), STAI (Laux et al., 1981), Becks Depression Inventory 2 (BDI-2) (Beck et al., 2011), and a questionnaire on the expectation of exercise in psychological and physiological domains. Furthermore, participants’ sleep quality, food intake, menstrual cycle phase, smoking, alcohol, and caffeine consumption on the study day and before the study days were assessed. The menstrual cycle phase was assessed by self-report on each experimental day where female participants indicated which phase applied to them (three phases: follicular, ovulatory, and luteal or hormonal contraceptives). Four female participants indicated they were on hormonal contraceptives. On both experimental days (Day 2 and Day 3) participants filled out the POMS before and after the study to monitor potential mood changes (results are reported in Table S22). Furthermore, participants were asked to fill out questionnaires including BDI, STAI, and a questionnaire concerning the side effects of NLX/SAL after both experimental days (Day 2 and Day 3; results are reported in Table S23). On the last experimental day (Day 3) participants had to indicate which pharmacological treatment they suspected to have received on which experimental day to capture potential expectations about the pharmacological treatment.

### Data Acquisition

#### Behavioural Data Acquisition

During the cycling blocks, heart rate, power output (Watts), and cadence (RPM) were recorded using MATLAB 2021b. For each cycling block the required power as well as the actual power maintained by the participant were displayed on a screen to allow for self-monitoring. During cycling, participants’ HR and SPO_2_ were constantly monitored and recorded. In one participant, the recording of the power output failed, and in five participants HR recordings in one session failed due to equipment issues. However, power output and HR maintained throughout cycling were visible and accessible at all times for the participant. Upon completion of the 10-minute cycling, participants were asked to rate their perceived exertion (RPE) on the BORG scale (no exertion (6) – maximal exertion (20)) (Borg, 1998) as well as to indicate their current mood (sad (0) – happy (10)). The transfer to the MR scanner commenced as quickly as possible (Time: *Mean* = 5 min, *SD* = 1 min) whilst maintaining the participants’ safety at all times. Inside the MR scanner, all stimulus presentation was realised using MATLAB R2016b and Psychophysics Toolbox (Version 3.0.19). Physiological data, including respiration and heart rate, was recorded at 1000 Hz with the Expression System (In Vivo, Gainesville, USA) using the spike2 software and a CED1401 system (Cambridge Electronic Design). A fixation cross and VAS scale were displayed on a screen inside the scanner bore. Thermal stimuli were applied using a TSA-2 Thermode with a 2.5×2.5 cm probe attached to the left volar forearm. Pressure stimuli were applied using a pressure cuff attached to the left upper arm and a CPAR outside the MR. The infusion pump was positioned outside the scanner with a ten-meter tube reaching into the scanner room to maintain a constant dose of NLX or SAL. After two pre-exposure stimuli of each modality, thermal and pressure stimuli were applied in an alternating fashion and at different calibrated intensities (randomised at 30, 50, 70 VAS). Each stimulus lasted 17 s in total (ramp-up, plateau 15 s, ramp-down) indicated by a visual cue (red fixation cross). Following each stimulus, the painfulness was rated on a VAS from ‘minimally painful’ (converted to 0 for analyses) to ‘almost unbearably painful’ (converted to 100 for analyses) by using a button box. The rating took place within 7 seconds after which an ITI of 7 seconds (jittered by 2 seconds) was introduced before commencing with the next trial. One MRI block consisted of nine thermal and nine pressure stimuli at three intensity levels (30, 50, 70 VAS). In total, participants spent 15 minutes per block inside the MR scanner.

#### MRI Data Acquisition

Functional magnetic resonance imaging (fMRI) data was acquired using a 3 Tesla Siemens system (Magnetom PRISMA; Siemens Healthcare, Erlangen, Germany) with a 64–channel head coil. Sixty slices were acquired (2 mm slice thickness) using T2*-weighted echo-planar imaging (EPI) with multiband factor 2 (repetition time (TR): 1.8 seconds; echo time (TE): 26ms; flip angle: 70°). The Field of View (FOV) was 240 mm and positioned to include the upper part of the medulla and the brainstem. The voxel size was thus 2 x 2 x 2 mm^3^. Four volumes at the beginning of each run were discarded. Furthermore, T1-weighted images were acquired for each subject using an MPRAGE sequence on the first experimental day (Day 2). Due to the nature of the study where participants cycled outside the MR scanner and had to be repositioned for each run, shimming and auto-align took place at the beginning of each run and before acquiring EPI images.

### Statistical Analyses

#### Behavioural Statistical Analyses

Behavioural statistical analyses were performed using MATLAB 2021b and RStudio (Version 2021.09.1). We used the *lmer* function from the *lme4* package (Version 1.1-35.1) (Bates et al., 2014) and the *lm* function from the *stats* package (Version 3.6.2) (*R: The R Project for Statistical Computing*) to conduct linear mixed effect (LMER) and linear models (LM) in R, respectively (Hox et al., 2010; Raudenbush and Bryk, 2001). See supplemental information (supplemental methods) for a detailed description of the statistical models used.

### MRI

#### Analyses Preprocessing

All MRI data was analysed using SPM12 (Wellcome Trust Centre for Neuroimaging, London, UK) and MATLAB 2021b (The Mathworks Inc. 2021b). The brain data was processed using a standardised SPM12-based pipeline (https://github.com/ChristianBuechel/spm_bids_pipeline.git) in BIDS format (Gorgolewski et al., 2016) to ensure data accessibility and reproducibility. The T1-weighted anatomical image was acquired on the first of the experimental days. The functional images from both experimental days were pre-processed together. After discarding the first four images of each run (dummy scans), the functional images were slice time corrected and realigned using rigid-body motion correction with six degrees of freedom. Next, a non-linear coregistration was performed using the mean EPI from the realignment and the T1 image. After this, the T1 image was normalised to MNI space (MNI152NLin2009cAsym as provided by CAT12 toolbox) using DARTEL. After this, transformation fields were created by combining the deformation field from the non-linear coregistration with the DARTEL flow fields to map the EPI images to T1 space and template space. A mask for the 1^st^-level GLMs was created based on the GM and WM from the T1 image and then warped into the EPI space and finally smoothed with a 3mm FWHM Gaussian kernel. After this, a second realignment took place but was constrained by a brain mask to minimise the effects of the eye movements. Finally, all EPI images were resliced in their individual space. Noise regressors for the first level analysis were created using six principal components of white matter responses and six principal components of CSF responses as well as a ROI at the posterior tip of the lateral ventricle (as used in Horing et al., 2019). As a final step, careful quality checks took place of all functional images generated after realignment, nonlinear co-registration, and normalisation through visual inspection, focusing especially on the successful alignment across both fMRI sessions to allow for subsequent analyses.

#### First-level analyses

The single-subject analyses of the brain data were performed in native space using a General Linear model (GLM) approach. The design matrices for each subject comprised eight blocks in total, each containing the stimulus onsets for heat and pressure stimuli at all three intensity levels (VAS 30, 50, 70) and a session constant. Additionally, the six motion parameters were augmented by their derivatives (12 parameters) and the squares of parameters and derivatives resulting in a total of 24 motion parameters (Friston et al., 1996). An additional spike correction was performed, where individual volumes with voxels with a deviation of 0.6 mm between each volume with its preceding volume of each run were flagged and individually modeled. In addition, we included RETROICOR (Glover et al., 2000) based physiological noise regressors to account for cardiac and respiratory-related motion (Brooks et al., 2008). This technique determines cardiac- and respiratory-related noise by allotting cardiac and respiratory phases to individual volumes within a time series. Subsequently, these assigned phases are utilised in a low-order Fourier expansion to generate time-course regressors elucidating signal variations attributed to cardiac or respiratory activities. A refined physiological noise model was applied, computing three cardiac and four respiratory harmonics, along with a multiplicative term incorporating interactions between cardiac and respiratory noise resulting in 18 regressors (Harvey et al., 2008). These resulting noise regressors were included in the first-level analyses. The contrast images from the first level analysis were then spatially normalised to MNI space using individual deformation fields (i.e. combining nonlinear coregistration and DARTEL spatial normalisation) and finally smoothed using a 6mm full width at half maximum (FWHM) Gaussian smoothing kernel. In addition to a hemodynamic response function (HRF) model, we also used a Finite Impulse Response model (FIR). This model was used to visualise the time course of BOLD responses after the onset of the pain stimulus. In this analysis, the stimulus duration was divided into 12 bins, each covering the duration of one TR spanning a total of 21.6 seconds after stimulus onset.

#### Second-level analyses

Subsequently, the spatially normalised and smoothed contrast images from the first-level analysis were used for random-effects group-level statistics. The successful application of heat pain was established by investigating the parametric contrast heat VAS 70 > 50 > 30, where higher stimulus intensities would recruit a more widespread (sub-)cortical pain network than lower intensities. To explore the effect of drug treatment in the brain, the contrast for the interaction of drug (SAL and NLX) and stimulus intensity (VAS 30, 50, 70) was calculated. Furthermore, we estimated the contrast heat NLX > heat SAL. To investigate the pain modulatory effect of exercise intensity in the SAL condition, the contrasts exercise high (SAL) > exercise low (SAL) and exercise low > high (SAL) were estimated. Furthermore, we extracted the parameter estimates from preregistered opioidergic ROIs (RVM, PAG, frontal midline) and estimated the LMER model corresponding to the behavioural analyses in the SAL condition. To confirm if NLX reduced the hypoalgesic effect, we estimated the contrasts interaction exercise intensity and drug treatment (positive and negative weight) as well as extracting the parameter estimates from the ROIs to conduct LMER models. To investigate the pain modulatory effect of exercise and fitness level in the SAL condition, the contrast exercise high (SAL) > exercise low (SAL) with the mean-centered covariate fitness level (weight-corrected FTP) was estimated. The parameter estimates from each condition were subtracted (LI exercise – HI exercise) and the difference in parameter estimates between both exercise conditions was visualised. To test the three-way interaction of fitness level, sex, and drug in the brain, a two-sample *t*-test of the contrast interaction exercise intensity and drug with the covariate fitness level has been conducted between males and females.

#### Correction for multiple comparisons

For whole-brain analyses, we used family-wise-error (FWE) rate as correction for multiple comparisons, and the according *P*-values are reported as *P_corr-WB_* = x.xx. Furthermore, we used a small volume correction mask (SVC) to correct for multiple comparisons, and the corrected *P*-values are reported as *P_corr-SVC_* = x.xx. This mask was based on the preregistered ROIs (rACC, vmPFC, PAG, RVM with an additional 1 mm FWHM Gaussian smoothing kernel (Figure S16 and Table S24). All thresholds for statistical significance were set to *P* < 0.05. Activation maps at an uncorrected level (*P_uncorr_* < 0.001) for each contrast are reported in the supplement.

## Acknowledgments

We thank Marilyn Mintah, Marie-Sophie Morgenroth, and Eileen Yawson for their support with data acquisition. We also thank Alexandra Tinnermann for her helpful comments. Further, we thank the radiographers at the Department of Systems Neuroscience and Jürgen Finsterbusch for providing the MR protocol.

## Author Contributions

Conceptualisation, C.B. and J.N.; Methodology, C.B. and J.N.; Investigation, C.B., J.N. and T.F.; Visualisation, J.N.; Writing – Original Draft, C.B. and J.N.; Funding Acquisition, C.B.; Resources, C.B.; Supervision, C.B.

## Ethics

Human subjects: All participants gave informed written consent. The study was approved by the Ethics Board of the Hamburg Medical Association (PV4932). We support inclusive, diverse, and equitable conduct of research.

## Additional Files

Supplemental Information: Supplemental Methods, Figures S1–S16, Tables S1-S24, and Supplemental References.

## Data availability

The datasets for the raw behavioural and fMRI data generated in the current study are available from the corresponding author on reasonable request and after publication. All necessary data to evaluate the results of the study are included in the manuscript and supplementary materials.

## Code availability

This study was programmed using MATLAB 2021b and Psychophysics Toolbox (Version 3.0.19). A custom fMRI preprocessing pipeline was used (https://github.com/ChristianBuechel/spm_bids_pipeline.git). The custom behavioural and fMRI analyses pipelines are available on the public repository https://github.com/jannenold/peep_analyses_mri_behavioural. For fMRI data visualisation, we used python-based software Nilearn (https://nilearn.github.io/dev/index.html#) and an MNI template with 1mm smoothing (MNI152_T1_1mm_brain).

## References

Allen H, Coggan A. 2012. Training and Racing with a Power Meter, 2nd Ed. VeloPress.

Allen H, Coggan A. 2006. Racing and training with a power meter. Boulder, CO: VeloPress.

Årnes AP, Nielsen CS, Stubhaug A, Fjeld MK, Hopstock LA, Horsch A, Johansen A, Morseth B, Wilsgaard T, Steingrímsdóttir ÓA. 2021. Physical activity and cold pain tolerance in the general population. Eur J Pain 25:637–650. doi:10.1002/ejp.1699

Årnes AP, Nielsen CS, Stubhaug A, Fjeld MK, Johansen A, Morseth B, Strand BH, Wilsgaard T, Steingrímsdóttir ÓA. 2023. Longitudinal relationships between habitual physical activity and pain tolerance in the general population. PLOS ONE 18:e0285041. doi:10.1371/journal.pone.0285041

Aslaksen PM, Bystad M, Vambheim SM, Flaten MA. 2011. Gender differences in placebo analgesia: event-related potentials and emotional modulation. Psychosom Med 73:193–199. doi:10.1097/PSY.0b013e3182080d73

Atlas LY, Wager TD. 2014. A Meta-analysis of Brain Mechanisms of Placebo Analgesia: Consistent Findings and Unanswered Questions. Handb Exp Pharmacol 225:37–69. doi:10.1007/978-3-662-44519-8_3

Bates D, Mächler M, Bolker B, Walker S. 2014. Fitting Linear Mixed-Effects Models using lme4.

Beck AT, Steer RA, Brown G. 2011. Beck Depression Inventory–II. doi:10.1037/t00742-000

Bodnar RJ, Kest B. 2010. Sex differences in opioid analgesia, hyperalgesia, tolerance and withdrawal: Central mechanisms of action and roles of gonadal hormones. Horm Behav, Sex and drugs: Sex differences and hormonal effects on drug abuse. 58:72–81. doi:10.1016/j.yhbeh.2009.09.012

Boecker H, Sprenger T, Spilker ME, Henriksen G, Koppenhoefer M, Wagner KJ, Valet M, Berthele A, Tolle TR. 2008. The Runner’s High: Opioidergic Mechanisms in the Human Brain. Cereb Cortex 18:2523–2531. doi:10.1093/cercor/bhn013

Borg G. 1998. Borg’s perceived exertion and pain scales, Borg’s perceived exertion and pain scales. Champaign, IL, US: Human Kinetics.

Borras MC, Becerra L, Ploghaus A, Gostic JM, DaSilva A, Gonzalez RG, Borsook D. 2004. FMRI Measurement of CNS Responses to Naloxone Infusion and Subsequent Mild Noxious Thermal Stimuli in Healthy Volunteers. J Neurophysiol 91:2723–2733. doi:10.1152/jn.00249.2003

Borszcz FK, Tramontin AF, Bossi AH, Carminatti LJ, Costa VP. 2018. Functional Threshold Power in Cyclists: Validity of the Concept and Physiological Responses. Int J Sports Med 39:737–742. doi:10.1055/s-0044-101546

Borszcz FK, Tramontin AF, Costa VP. 2020. Reliability of the Functional Threshold Power in Competitive Cyclists. Int J Sports Med 41:175–181. doi:10.1055/a-1018-1965

Brellenthin AG, Crombie KM, Cook DB, Sehgal N, Koltyn KF. 2017. Psychosocial Influences on Exercise-Induced Hypoalgesia. Pain Med 18:538–550. doi:10.1093/pm/pnw275

Brooks JCW, Beckmann CF, Miller KL, Wise RG, Porro CA, Tracey I, Jenkinson M. 2008. Physiological noise modelling for spinal functional magnetic resonance imaging studies. NeuroImage 39:680–692. doi:10.1016/j.neuroimage.2007.09.018

Brooks JCW, Davies W-E, Pickering AE. 2017. Resolving the Brainstem Contributions to Attentional Analgesia. J Neurosci 37:2279–2291. doi:10.1523/JNEUROSCI.2193-16.2016

Bulls HW, Freeman EL, Anderson AJ, Robbins MT, Ness TJ, Goodin BR. 2015. Sex differences in experimental measures of pain sensitivity and endogenous pain inhibition. J Pain Res 8:311–320. doi:10.2147/JPR.S84607

Coghill RC. 2020. The Distributed Nociceptive System: A Framework for Understanding Pain. Trends Neurosci 43:780–794. doi:10.1016/j.tins.2020.07.004

Corder G, Castro DC, Bruchas MR, Scherrer G. 2018. Endogenous and Exogenous Opioids in Pain. Annu Rev Neurosci 41:453–473. doi:10.1146/annurev-neuro-080317-061522

Crombie KM, Brellenthin AG, Hillard CJ, Koltyn KF. 2018. Endocannabinoid and opioid system interactions in exercise-induced hypoalgesia. Pain Med 19:118–123. doi:10.1093/pm/pnx058

Curran SL, Andrykowski MA, Studts JL. 1995. Short Form of the Profile of Mood States (POMS-SF): Psychometric information. Psychol Assess 7:80–83. doi:10.1037/1040-3590.7.1.80

Denham J, Scott-Hamilton J, Hagstrom AD, Gray AJ. 2020. Cycling Power Outputs Predict Functional Threshold Power and Maximum Oxygen Uptake. J Strength Cond Res 34:3489–3497. doi:10.1519/JSC.0000000000002253

Desai S, Borg B, Cuttler C, Crombie KM, Rabinak CA, Hill MN, Marusak HA. 2022. A Systematic Review and Meta-Analysis on the Effects of Exercise on the Endocannabinoid System. Cannabis Cannabinoid Res 7:388–408. doi:10.1089/can.2021.0113

Droste C, Greenlee MW, Schreck M, Roskamm H. 1991. Experimental pain thresholds and plasma beta-endorphin levels during exercise. Med Sci Sports Exerc 23:334.

Droste C, Meyer-Blankenburg H, Greenlee MW, Roskamm H. 1988. Effect of physical exercise on pain thresholds and plasma beta-endorphins in patients with silent and symptomatic myocardial ischaemia. Eur Heart J 9 Suppl N:25–33. doi:10.1093/eurheartj/9.suppl_n.25

Eippert F, Bingel U, Schoell ED, Yacubian J, Klinger R, Lorenz J, Büchel C. 2009. Activation of the Opioidergic Descending Pain Control System Underlies Placebo Analgesia. Neuron 63:533–543. doi:10.1016/j.neuron.2009.07.014

Ellingson LD, Stegner AJ, Schwabacher IJ, Koltyn KF, Cook DB. 2016. Exercise Strengthens Central Nervous System Modulation of Pain in Fibromyalgia. Brain Sci 6:8. doi:10.3390/brainsci6010008

Friston KJ, Williams S, Howard R, Frackowiak RSJ, Turner R. 1996. Movement-Related effects in fMRI time-series. Magn Reson Med 35:346–355. doi:10.1002/mrm.1910350312

Fuss J, Steinle J, Bindila L, Auer MK, Kirchherr H, Lutz B, Gass P. 2015. A runner’s high depends on cannabinoid receptors in mice. Proc Natl Acad Sci 112:13105–13108. doi:10.1073/pnas.1514996112

Ge H-Y, Madeleine P, Arendt-Nielsen L. 2004. Sex differences in temporal characteristics of descending inhibitory control: an evaluation using repeated bilateral experimental induction of muscle pain. PAIN 110:72. doi:10.1016/j.pain.2004.03.005

Geisler M, Eichelkraut L, Miltner WHR, Weiss T. 2019. An fMRI study on runner’s high and exercise-induced hypoalgesia after a 2-h-run in trained non-elite male athletes. Sport Sci Health. doi:10.1007/s11332-019-00592-8

Geva N, Defrin R. 2013. Enhanced pain modulation among triathletes: A possible explanation for their exceptional capabilities. PAIN® 154:2317–2323. doi:10.1016/j.pain.2013.06.031

Glover GH, Li T-Q, Ress D. 2000. Image-based method for retrospective correction of physiological motion effects in fMRI: RETROICOR. Magn Reson Med 44:162–167. doi:10.1002/1522-2594(200007)44:1<162::AID-MRM23>3.0.CO;2-E

Gomolka S, Vaegter HB, Nijs J, Meeus M, Gajsar H, Hasenbring MI, Titze C. 2019. Assessing Endogenous Pain Inhibition: Test-Retest Reliability of Exercise-Induced Hypoalgesia in Local and Remote Body Parts after Aerobic Cycling. Pain Med 20:2272–2282. doi:10.1093/pm/pnz131

Gorgolewski KJ, Auer T, Calhoun VD, Craddock RC, Das S, Duff EP, Flandin G, Ghosh SS, Glatard T, Halchenko YO, Handwerker DA, Hanke M, Keator D, Li X, Michael Z, Maumet C, Nichols BN, Nichols TE, Pellman J, Poline J-B, Rokem A, Schaefer G, Sochat V, Triplett W, Turner JA, Varoquaux G, Poldrack RA. 2016. The brain imaging data structure, a format for organizing and describing outputs of neuroimaging experiments. Sci Data 3:160044. doi:10.1038/sdata.2016.44

Granot M, Weissman-Fogel I, Crispel Y, Pud D, Granovsky Y, Sprecher E, Yarnitsky D. 2008. Determinants of endogenous analgesia magnitude in a diffuse noxious inhibitory control (DNIC) paradigm: Do conditioning stimulus painfulness, gender and personality variables matter? Pain 136:142–149. doi:10.1016/j.pain.2007.06.029

Graven-Nielsen T, Vaegter HB, Finocchietti S, Handberg G, Arendt-Nielsen L. 2015. Assessment of musculoskeletal pain sensitivity and temporal summation by cuff pressure algometry: a reliability study. Pain 156:2193– 2202. doi:10.1097/j.pain.0000000000000294

Gurevich M, Kohn PM, Davis C. 1994. Exercise-induced analgesia and the role of reactivity in pain sensitivity. J Sports Sci 12:549–559. doi:10.1080/02640419408732205

Haier RJ, Quaid K, Mills JSC. 1981. Naloxone Alters Pain Perception After Jogging. Psychiatry Res 5:231–232. doi:10.1016/0165-1781(81)90052-4

Harvey AK, Pattinson KTS, Brooks JCW, Mayhew SD, Jenkinson M, Wise RG. 2008. Brainstem functional magnetic resonance imaging: Disentangling signal from physiological noise. J Magn Reson Imaging 28:1337–1344. doi:10.1002/jmri.21623

Horing B, Sprenger C, Büchel C. 2019. The parietal operculum preferentially encodes heat pain and not salience. PLoS Biol 17:e3000205. doi:10.1371/journal.pbio.3000205

Hox JJ, Moerbeek M, Schoot R van de. 2010. Multilevel Analysis: Techniques and Applications, Second Edition, 2nd ed. New York: Routledge.

Janal MN, Colt EWD, Crawford W, Glusman M. 1984. Pain Sensitivity, Mood and Plasma Endocrine Levels in Man Following Long-Distance Running: Effects of Naloxone. Pain 19:13–25. doi:10.1016/0304-3959(84)90061-7

Jones AM, Burnley M, Black MI, Poole DC, Vanhatalo A. 2019. The maximal metabolic steady state: redefining the ‘gold standard.’ Physiol Rep 7:e14098. doi:10.14814/phy2.14098

Jones MD, Nuzzo JL, Taylor JL, Barry BK. 2019. Aerobic Exercise Reduces Pressure More Than Heat Pain Sensitivity in Healthy Adults. Pain Med 20:1534–1546. doi:10.1093/pm/pny289

Jones MD, Valenzuela T, Booth J, Taylor JL, Barry BK. 2017. Explicit Education About Exercise-Induced Hypoalgesia Influences Pain Responses to Acute Exercise in Healthy Adults: A Randomized Controlled Trial. J Pain 18:1409–1416. doi:10.1016/j.jpain.2017.07.006

Klich S, Krymski I, Michalik K, Kawczyński A. 2018. Effect of short-term cold-water immersion on muscle pain sensitivity in elite track cyclists. Phys Ther Sport Off J Assoc Chart Physiother Sports Med 32:42–47. doi:10.1016/j.ptsp.2018.04.022

Koltyn KF. 2002. Exercise-Induced Hypoalgesia and Intensity of Exercise. Sports Med 32:477–487. doi:10.2165/00007256-200232080-00001

Koltyn KF. 2000. Analgesia Following Exercise. Sports Med 29:85–98. doi:10.2165/00007256-200029020-00002

Koltyn KF, Brellenthin AG, Cook DB, Sehgal N, Hillard C. 2014. Mechanisms of exercise-induced hypoalgesia. J Pain 15:1294–1304. doi:10.1016/j.jpain.2014.09.006

Koltyn KF, Garvin AW, Gardiner RL, Nelson TF. 1996. Perception of pain following aerobic exercise. Med Sci Sports Exerc 28:1418–1421.

Koltyn KF, Knauf M t., Brellenthin A g. 2013. Temporal summation of heat pain modulated by isometric exercise. Eur J Pain 17:1005–1011. doi:10.1002/j.1532-2149.2012.00264.x

Kragel PA, Kano M, Van Oudenhove L, Ly HG, Dupont P, Rubio A, Delon-Martin C, Bonaz BL, Manuck SB, Gianaros PJ, Ceko M, Reynolds Losin EA, Woo C-W, Nichols TE, Wager TD. 2018. Generalizable representations of pain, cognitive control, and negative emotion in medial frontal cortex. Nat Neurosci 21:283–289. doi:10.1038/s41593-017-0051-7

Kut E, Candia V, Overbeck J von, Pok J, Fink D, Folkers G. 2011. Pleasure-Related Analgesia Activates Opioid- Insensitive Circuits. J Neurosci 31:4148–4153. doi:10.1523/JNEUROSCI.3736-10.2011

Laux L, Glanzmann P, Schaffner P, Spielberger CD. 1981. Das State-Trait Angstinventar. Theoretische Grundlagen und Handanweisung.

Leknes S, Atlas LY. 2020. Flawed methodology undermines conclusions about opioid-induced pleasure: implications for psychopharmacology. Br J Anaesth 124:e29–e33. doi:10.1016/j.bja.2019.10.006

Lesnak JB, Sluka KA. 2020. Mechanism of exercise-induced analgesia: what we can learn from physically active animals. PAIN Rep 5:e850. doi:10.1097/PR9.0000000000000850

Lindheimer JB, Szabo A, Raglin JS, Beedie C, Carmichael KE, O’Connor PJ. 2020. Reconceptualizing the measurement of expectations to better understand placebo and nocebo effects in psychological responses to exercise. Eur J Sport Sci 20:338–346. doi:10.1080/17461391.2019.1674926

Mackey J, Horner K. 2021. What is known about the FTP ^20^ test related to cycling? A scoping review. J Sports Sci 39:2735–2745. doi:10.1080/02640414.2021.1955515

Martin LJ, Acland EL, Cho C, Gandhi W, Chen D, Corley E, Kadoura B, Levy T, Mirali S, Tohyama S, Khan S, MacIntyre LC, Carlson EN, Schweinhardt P, Mogil JS. 2019. Male-Specific Conditioned Pain Hypersensitivity in Mice and Humans. Curr Biol 29:192–201.e4. doi:10.1016/j.cub.2018.11.030

Mc Grath E, Mahony N, Fleming N, Donne B. 2019. Is the FTP Test a Reliable, Reproducible and Functional Assessment Tool in Highly-Trained Athletes? Int J Exerc Sci 12:1334–1345.

Micalos PS, Arendt-Nielsen L. 2016. Differential pain response at local and remote muscle sites following aerobic cycling exercise at mild and moderate intensity. SpringerPlus 5:1–5. doi:10.1186/s40064-016-1721-8

Mogil JS. 2020. Qualitative sex differences in pain processing: emerging evidence of a biased literature. Nat Rev Neurosci 21:353–365. doi:10.1038/s41583-020-0310-6

Mogil JS. 2012. Sex differences in pain and pain inhibition: multiple explanations of a controversial phenomenon. Nat Rev Neurosci 13:859–866. doi:10.1038/nrn3360

Mogil JS, Chanda ML. 2005. The case for the inclusion of female subjects in basic science studies of pain. PAIN 117:1. doi:10.1016/j.pain.2005.06.020

Mumford J. 2012. A power calculation guide for FMRI studies. Soc Cogn Affect Neurosci 7:738–42. doi:10.1093/scan/nss059

Naugle KM, Fillingim RB, Riley JL. 2012. A meta-analytic review of the hypoalgesic effects of exercise. J Pain 13:1139–1150. doi:10.1016/j.jpain.2012.09.006

Naugle KM, Naugle KE, Fillingim RB, Samuels B, Riley JLI. 2014. Intensity Thresholds for Aerobic Exercise- Induced Hypoalgesia. Med Sci Sports Exerc 46:817–825. doi:10.1249/MSS.0000000000000143

Naugle KM, Riley JL. 2014. Self-reported Physical Activity Predicts Pain Inhibitory and Facilitatory Function. Med Sci Sports Exerc 46:622–629. doi:10.1249/MSS.0b013e3182a69cf1

Niesters M, Dahan A, Kest B, Zacny J, Stijnen T, Aarts L, Sarton E. 2010. Do sex differences exist in opioid analgesia? A systematic review and meta-analysis of human experimental and clinical studies. PAIN 151:61. doi:10.1016/j.pain.2010.06.012

Olausson B, Eriksson E, Ellmarker L, Rydenhag B, Shyu B-C, Andersson SA. 1986. Effects of naloxone on dental pain threshold following muscle exercise and low frequency transcutaneous nerve stimulation: a comparative study in man. Acta Physiol Scand 126:299–305. doi:10.1111/j.1748-1716.1986.tb07818.x

Padawer WJ, Levine FM. 1992. Exercise-induced analgesia: fact or artifact? Pain 48:131–135. doi:10.1016/0304-3959(92)90048-G

Petrovic P, Kalso E, Petersson KM, Ingvar M. 2002. Placebo and Opioid Analgesia-- Imaging a Shared Neuronal Network. Science 295:1737–1740. doi:10.1126/science.1067176

Raudenbush SW, Bryk AS. 2001. Hierarchical Linear Models: Applications and Data Analysis Methods, 2nd Edition. ed. Thousand Oaks: SAGE Publications, Inc.

Ruble SB, Hoffman MD, Shepanski MA, Valic Z, Buckwalter JB, Clifford PS. 2005. Thermal Pain Perception After Aerobic Exercise. Arch Phys Med Rehabil 86:1019–1023. doi:10.1016/j.apmr.2004.09.024

Saanijoki T, Kantonen T, Pekkarinen L, Kalliokoski K, Hirvonen J, Malén T, Tuominen L, Tuulari JJ, Arponen E, Nuutila P, Nummenmaa L. 2022. Aerobic Fitness is Associated with Cerebral mu-Opioid Receptor Activation in Healthy Humans. Med Sci Sports Exerc. doi:10.1249/MSS.0000000000002895

Saanijoki T, Tuominen L, Tuulari JJ, Nummenmaa L, Arponen E, Kalliokoski K, Hirvonen J. 2018. Opioid Release after High-Intensity Interval Training in Healthy Human Subjects. Neuropsychopharmacology 43:246–254. doi:10.1038/npp.2017.148

Scheef L, Jankowski J, Daamen M, Weyer G, Klingenberg M, Renner J, Mueckter S, Schürmann B, Musshoff F, Wagner M, Schild HH, Zimmer A, Boecker H. 2012. An fMRI study on the acute effects of exercise on pain processing in trained athletes. Pain 153:1702–1714. doi:10.1016/j.pain.2012.05.008

Schmitt A, Wallat D, Stangier C, Martin JA, Schlesinger-Irsch U, Boecker H. 2020. Effects of fitness level and exercise intensity on pain and mood responses. Eur J Pain 24:568–579. doi:10.1002/ejp.1508

Schoell ED, Bingel U, Eippert F, Yacubian J, Christiansen K, Andresen H, May A, Buechel C. 2010. The Effect of Opioid Receptor Blockade on the Neural Processing of Thermal Stimuli. PLoS ONE 5:e12344. doi:10.1371/journal.pone.0012344

Segerdahl AR, Mezue M, Okell TW, Farrar JT, Tracey I. 2015. The dorsal posterior insula subserves a fundamental role in human pain. Nat Neurosci 18:499–500. doi:10.1038/nn.3969

Shackman AJ, Salomons TV, Slagter HA, Fox AS, Winter JJ, Davidson RJ. 2011. The integration of negative affect, pain and cognitive control in the cingulate cortex. Nat Rev Neurosci 12:154–167. doi:10.1038/nrn2994

Siebers M, Biedermann SV, Bindila L, Lutz B, Fuss J. 2021. Exercise-induced euphoria and anxiolysis do not depend on endogenous opioids in humans. Psychoneuroendocrinology 126:105173. doi:10.1016/j.psyneuen.2021.105173

Sluka KA, Frey-Law L, Hoeger Bement M. 2018. Exercise-induced pain and analgesia? Underlying mechanisms and clinical translation. Pain 159:S91–S97. doi:10.1097/j.pain.0000000000001235

Smith LD. 2004. The effects of competition and exercise on pain perception.

Sørensen A, Aune TK, Rangul V, Dalen T. 2019. The Validity of Functional Threshold Power and Maximal Oxygen Uptake for Cycling Performance in Moderately Trained Cyclists. Sports 7:217. doi:10.3390/sports7100217

Sprenger C, Eippert F, Finsterbusch J, Bingel U, Rose M, Büchel C. 2012. Attention Modulates Spinal Cord Responses to Pain. Curr Biol 22:1019–1022. doi:10.1016/j.cub.2012.04.006

Stagg NJ, Mata HP, Ibrahim MM, Henriksen EJ, Porreca F, Vanderah TW, Malan TP. 2011. Regular exercise reverses sensory hypersensitivity in a rat neuropathic model: role of endogenous opioids. Anesthesiology 114:940–948. doi:10.1097/ALN.0b013e318210f880

Sternberg WF, Boka C, Kas L, Alboyadjia A, Gracely RH. 2001. Sex-Dependent Components of the Analgesia Produced by Athletic Competition. J Pain 2:65–74. doi:10.1054/jpai.2001.18236

Tesarz J, Schuster AK, Hartmann M, Gerhardt A, Eich W. 2012. Pain perception in athletes compared to normally active controls: a systematic review with meta-analysis. Pain 153:1253–1262. doi:10.1016/j.pain.2012.03.005

Tinnermann A, Sprenger C, Büchel C. 2022. Opioid analgesia alters corticospinal coupling along the descending pain system in healthy participants. eLife. doi:10.7554/eLife.74293

Tracey I, Mantyh PW. 2007. The Cerebral Signature for Pain Perception and Its Modulation. Neuron 55:377–391. doi:10.1016/j.neuron.2007.07.012

Tran TD, Wang H, Tandon A, Hernandez-Garica L, Casey KL. 2010. Temporal summation of heat pain in humans: Evidence supporting thalamocortical modulation. Pain 150:93–102. doi:10.1016/j.pain.2010.04.001

Trøstheim M, Eikemo M, Haaker J, Frost JJ, Leknes S. 2022. Opioid antagonism in humans: a primer on optimal dose and timing for central mu-opioid receptor blockade. Neuropsychopharmacology 1–9. doi:10.1038/s41386-022-01416-z

Vaegter H, Handberg G, Jorgensen M, Kinly A, Graven-Nielsen T. 2015. Aerobic exercise and cold pressor test induce hypoalgesia in active and inactive men and women. Pain Med. doi:10.1111/pme.12641

Vaegter HB, Bjerregaard LK, Redin M-M, Rasmussen SH, Graven-Nielsen T. 2019. Hypoalgesia after bicycling at lactate threshold is reliable between sessions. Eur J Appl Physiol 119:91–102. doi:10.1007/s00421-018-4002-0

Vaegter HB, Handberg G, Graven-Nielsen T. 2014. Similarities between exercise-induced hypoalgesia and conditioned pain modulation in humans. Pain 155:158–167. doi:10.1016/j.pain.2013.09.023

Vaegter HB, Jones MD. 2020. Exercise-induced hypoalgesia after acute and regular exercise: experimental and clinical manifestations and possible mechanisms in individuals with and without pain. Pain Rep 5:e823. doi:10.1097/PR9.0000000000000823

Wewege MA, Jones MD. 2020. Exercise-Induced Hypoalgesia in Healthy Individuals and People With Chronic

Wiech K, Lin C, Brodersen KH, Bingel U, Ploner M, Tracey I. 2010. Anterior Insula Integrates Information about Salience into Perceptual Decisions about Pain. J Neurosci 30:16324–16331. doi:10.1523/JNEUROSCI.2087-10.2010

Wilson EB. 1927. Probable Inference, the Law of Succession, and Statistical Inference. J Am Stat Assoc 22:209–212. doi:10.1080/01621459.1927.10502953

Wong S, Burnley M, Mauger A, Fenghua S, Hopker J. 2022. Functional threshold power is not a valid marker of the maximal metabolic steady state. J Sports Sci 40:2578–2584. doi:10.1080/02640414.2023.2176045

World Medical Association. 2013. World Medical Association Declaration of Helsinki: ethical principles for medical research involving human subjects. JAMA 310:2191–2194. doi:10.1001/jama.2013.281053

Wu B, Zhou L, Chen C, Wang J, Hu L, Wang X. 2022. Effects of Exercise-induced Hypoalgesia and Its Neural Mechanisms. Med Sci Sports Exerc 54:220–231. doi:10.1249/MSS.0000000000002781

