## Supplemental Materials for "Acute aerobic exercise intensity does not modulate pain potentially due to differences in fitness levels and sex effects – results from a pharmacological fMRI study"

**Table of Contents**

Behavioural Statistical Analyses ........................................................................................................................5

Figure S1. Effect of exercise intensity, drug treatment, and sex on pressure pain modulation. (A) Pressure pain ratings pooled across all stimulus intensities in the SAL (blue) and NLX (orange) conditions at low and high exercise intensity. There was no hypoalgesic effect evident in the behavioural pain ratings comparing HI to LI exercise in the SAL condition ($\beta$ = 0.57, *CI* [-1.73, 2.86], *SE* = 1.17, $t$(1354) = 0.48, $P$ = 0.63; blue bars) as well as no interaction of drug treatment and exercise intensity on pressure pain ratings ($\beta$ = -1.43, *CI* [-4.87, 2.01], *SE* = 1.75, $t$(2756.02) = -0.82, $P$ = 0.42). Bars depict the mean ratings in the SAL and NLX conditions at both exercise intensities averaged across all stimulus intensities. Individual data points depict subject-specific mean pain ratings. Error bars depict the SEM ($N$ = 39). (B) Subject-specific differences in pressure pain ratings (dots) between low-intensity (LI) and high-intensity (HI) exercise conditions (LI – HI exercise pain ratings) and corresponding regression line pooled across all stimulus intensities in the SAL condition. Fitness level (FTP) showed no significant relation to pressure pain ratings ($r$ = 0.25, $P$ = 0.13) and no significant main effect of FTP ($\beta$ = 3.16, *CI* [-1.64, 7.97], *SE* = 2.37, $t$(38) = 1.34, $P$ = 0.19) on difference ratings. (C-D) No significant interaction of drug treatment, exercise intensity, and sex on difference pain ratings ($\beta$ =-7.97, *CI* [-18.67, 2.73], *SE* = 5.51, $t$(190) = -1.45, $P$ = 0.15 with exercise-induced pain modulation in the (C) SAL and (D) NLX condition showing no significant difference between males (red) and females (blue). In the NLX condition males show a trend where with increasing fitness levels (FTP) the hypoalgesic response diminished. ........................................................................................................................7

Figure S2. Uncorrected activation map for parametric contrast (VAS 70 > 50 > 30) of heat pain in saline (SAL) condition ($P$ < 0.001). BOLD activation at $P$ < 0.001 uncorrected superimposed onto mean T1 along 134 slices (2 slice steps) in the z-axis. ........................................................................................................................8

Figure S3. Expectation of acute exercise on pain. (A) Ratings for acute exercise on different pain types and ranging between -3 (‘large decrease’) and 3 (‘large increase’) with 0 denoting ‘no effect’. There was no significant effect for muscle pain ($t$(38) = 1.78, $P$ = 0.08, $M$ = 0.39, *SE* = 0.12), joint pain ($t$(38) = -0.12, $P$ = 0.90, $M$ = -0.03, *SE* = 0.11), or ‘whole-body pain ($t$(38) = -1.05, $P$ = 0.30, $M$ = -0.21, *SE* = 0.12) suggesting there to be no expectation effect on these pain domains in the overall sample. Boxplots depict the distribution of the data with dots depicting subject-specific expectation ratings. Black dots depict the mean expectation ratings averaged across subjects and error bars depict the SEM. (B) Correlation between difference pain ratings (LI-HI exercise intensity) in the saline (SAL) condition and expectation ratings with each facet depicting a different pain type for each subject (dot). This analysis yielded no significant correlation in either of the pain domains (joint pain: $r$ = 0.11, $P$ = 0.49; muscle pain: $r$ = -0.07, $P$ = 0.68; whole-body pain: $r$ = 0.07, $P$ = 0.68). Regression lines are visualised and shaded areas represent the SEM ($N$ = 39). ....................................9

Figure S4. Uncorrected activation map for contrast interaction stimulus intensity x drug treatment ($P$ < 0.001). BOLD activation at $P$ < 0.001 uncorrected superimposed onto mean T1 along 134 slices (2 slice steps) in the z-axis. ........................................................................................................................10

Figure S5. Cortical drug treatment effect in the anterior Insula. (A) Activation for contrast: heat NLX > heat SAL in right anterior insula ($MNI_{xyz}$: 46, 8, 6; T = 5.70, $P_{corr\text{-}WB}$ = 0.08) superimposed onto MNI template brain along x and z dimension. (B) Parameter estimates for saline (blue) and naloxone (orange) conditions for the respective peak voxel in the right anterior insula ($MNI_{xyz}$: 46, 8, 6). Individual dots indicate subject-specific mean parameter estimates whereas solid dots indicate the overall mean for drug treatment conditions at stimulus intensity. (C) Difference between parameter estimates of the respective peak voxels between naloxone and saline condition at each stimulus intensity. (D) Time course of BOLD response for saline (blue) and naloxone (orange) condition for heat pain at VAS 70 in the respective peak voxels. The shaded areas around the curves represent the standard error of the mean ($n$ = 39). The grey solid lines indicate stimulus start and stimulus end and the shaded grey area displays the approximate time window for BOLD response (5 seconds after stimulus onset). ........................................................................................................................11

Figure S6. Uncorrected activation map for contrast exercise high > low of heat pain in saline (SAL) condition ($P < 0.01$). BOLD activation at $P < 0.01$ uncorrected superimposed onto mean T1 along 134 slices (2 slice steps) in the z-axis. ....12
Figure S7. Uncorrected activation map for contrast exercise low > high of heat pain in saline (SAL) condition ($P < 0.01$). BOLD activation at $P < 0.01$ uncorrected superimposed onto mean T1 along 134 slices (2 slice steps) in the z-axis. ....13
Figure S8. Effect of exercise intensity and drug treatment on heat pain ratings at different stimulus intensities (VAS 30, 50, 70). Bars depict the mean ratings in the saline (SAL; blue) and naloxone (NLX; orange) conditions. Individual data points depict subject-specific mean pain ratings. Error bars depict the SEM. The LMER was extended to include the stimulus intensity and yielded a significant main effect of stimulus intensity ($\beta$ = 1.39, *CI* [1.31, 1.47], *SE* = 0.04, $t(2753.12) = -34.082$, $P < 0.001$) and a significant interaction of stimulus intensity and drug treatment ($\beta$ = 0.12, *CI* [0.01, 0.24], *SE* = 0.06, $t(2751) = 2.13$, $P = 0.03$), but no significant interaction of exercise intensity, drug treatment, and stimulus intensity (β = -0.05, CI [-0.20, 0.11], SE = 0.08, t(2751) = -0.56, P = 0.58). ....14
Figure S9. Uncorrected activation map for contrast interaction exercise intensity and drug treatment (pos) of heat pain ($P < 0.01$). BOLD activation at $P < 0.01$ uncorrected superimposed onto mean T1 along 134 slices (2 slice steps) in the z-axis. ....15
Figure S10. Uncorrected activation map for contrast interaction exercise intensity and drug treatment (neg) of heat pain ($P < 0.01$). BOLD activation at $P < 0.01$ uncorrected superimposed onto mean T1 along 134 slices (2 slice steps) in the z-axis. ....16
Figure S11. Effect of exercise intensity and fitness level on absolute pain ratings and medial frontal cortex (mFC) activation. (A) Subject-specific heat pain ratings (dots) between low-intensity (green) and high-intensity (purple) exercise conditions and corresponding regression lines pooled across all stimulus intensities in the saline condition. (B) Cortical activation for contrast: exercise high > exercise low with mean-centered covariate FTP (weight-corrected) in right mFC ($MNI_{xyz}$: 6, 45, 10; T = 4.59, $P_{corr\text{-}SVC} = 0.05$) across all stimulus intensities in the saline condition superimposed onto the MNI template brain. (C) Parameter estimates of low-intensity (LI; green) and high-intensity (HI; purple) exercise conditions from respective peak voxel plotted for each subject depending on fitness levels (FTP) and pooled across stimulus intensities. ....17
Figure S16. Small Volume Correction mask based on preregistered ROIs. Small volume correction mask with 1 mm smoothing in MNI space superimposed onto MNI Template with slices along x (top) and z-axis (bottom). Colour codes are according to ROI: red = rostral ventral medulla (RVM); yellow = Frontal Midline (comprises anterior cingulate cortex (ACC) and ventromedial prefrontal cortex (vmPFC), green = PAG. ..22

**Behavioural Statistical Analyses**

For behavioral analyses, pain ratings of the VAS with the endpoints ‘minimally painful’ and ‘almost unbearably painful’ were converted to a numerical scale with the endpoints 0 and 100, respectively. In every LMER model, heat pain ratings or differential heat pain ratings (LI-HI pain ratings) served as the dependent variable, the subject as well as the number of pain ratings (1-9 per block) were included as random effects, whereas the order of drug treatment administration (naloxone or saline first) was included as a fixed effect. In the models that included pain ratings from all stimulus intensities (VAS 30, 50, 70), the stimulus intensity was also included as a fixed effect to account for the variance. Post-hoc paired samples *t*-tests (two-sided) were calculated on the LMER models using the R package *emmeans* (Version 1.10.0). The Tukey Method was used for adjusting the resulting *p*-values as implemented in the *emmeans* package. In the instances where the LM was used, heat pain ratings or differential heat pain ratings (LI-HI pain ratings) served as the dependent variable, and order of drug administration was included as an additional independent variable.

To verify the successful application of heat pain, the respective pain ratings in the saline condition only were used as the dependent variable in the LMER model and stimulus intensity (VAS 30, 50, 70) served as a fixed effect. The successful exercise intervention was measured by the average power output, heart rate, and rating of perceived exertion. The power output (watt) and heart rate (bpm) were measured throughout the cycling blocks. The ratings of perceived exertion were measured after each cycling block and a mean RPE value was calculated across the low and high exercise conditions. Paired *t*-tests (two-sided) were conducted for the mean power output, heart rate, and RPE between the low and high exercise intensity conditions. Cohens *d* has been calculated as effect size for within-subject samples. To verify the successful drug administration, we conducted LMER models for heat pain ratings with drug and stimulus intensity as well as their interaction as fixed effects. The difference between pain ratings between both drug treatment conditions has been calculated and an LMER model with stimulus intensity as a fixed effect has been calculated. Concerning the potentially diverging effect of drug treatment depending on sex, we conducted an LMER model with drug and stimulus intensity on heat pain ratings for males and females separately. Again, the difference between pain ratings between both drug treatment conditions has been calculated and an LMER model with stimulus intensity as a fixed effect has been calculated for both sexes.

To test our hypothesis of exercise-induced pain modulation, an LMER model with the fixed effects of exercise intensity on heat pain ratings has been calculated. To capture the effect more reliably, the heat pain ratings after high-intensity exercise were subtracted from heat pain ratings after low-intensity exercise. Positive difference scores would indicate hypoalgesia whereas a negative score would indicate hyperalgesia following high-intensity exercise. In a LM, the fitness level (weight-corrected FTP) served as an independent variable, and the differential heat pain ratings as the dependent variable. To model the interaction of fitness level, sex, and drug the differential pain ratings served as the dependent variable for the LMER model, the fitness levels (weight-corrected FTP), drug, and sex as well as the three-way interaction served as fixed effects.

The mood ratings provided on the POMS on each experimental day pre and post-measurements were summarised in four domains (dejection, fatigue, discontent, drive) and analysed using Wilcoxon signed rank test (Wilcoxon, 1945). There were no significant differences in the post-treatment measurements between the drug treatment conditions (Table S22). Furthermore, potential differences in side effects were analysed between the drug treatment conditions using the Wilcoxon signed rank test (Wilcoxon, 1945) and showed no significant differences between the drug treatment conditions (Table S23).

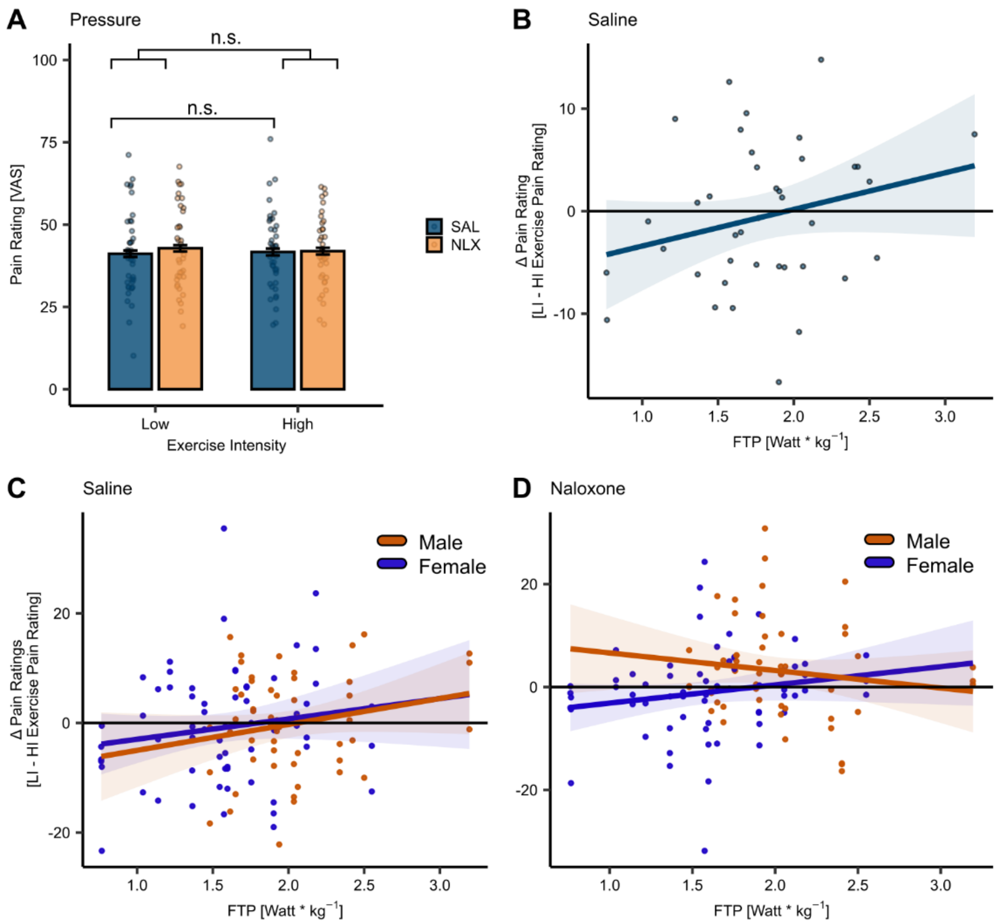

**Figure S1**. Effect of exercise intensity, drug treatment, and sex on pressure pain modulation. (A) Pressure pain ratings pooled across all stimulus intensities in the SAL (blue) and NLX (orange) conditions at low and high exercise intensity. There was no hypoalgesic effect evident in the behavioural pain ratings comparing HI to LI exercise in the SAL condition ($\beta$ = 0.57, *CI* [-1.73, 2.86], *SE* = 1.17, *t*(1354) = 0.48, *P* = 0.63; blue bars) as well as no interaction of drug treatment and exercise intensity on pressure pain ratings ($\beta$ = -1.43, *CI* [-4.87, 2.01], *SE* = 1.75, *t*(2756.02) = -0.82, *P* = 0.42). Bars depict the mean ratings in the SAL and NLX conditions at both exercise intensities averaged across all stimulus intensities. Individual data points depict subject-specific mean pain ratings. Error bars depict the SEM (*N* = 39). (B) Subject-specific differences in pressure pain ratings (dots) between low-intensity (LI) and high-intensity (HI) exercise conditions (LI – HI exercise pain ratings) and corresponding regression line pooled across all stimulus intensities in the SAL condition. Fitness level (FTP) showed no significant relation to pressure pain ratings (*r* = 0.25, *P* = 0.13) and no significant main effect of FTP ($\beta$ = 3.16, *CI* [-1.64, 7.97], *SE* = 2.37, *t*(38) = 1.34, *P* = 0.19) on difference ratings. (C-D) No significant interaction of drug treatment, exercise intensity, and sex on difference pain ratings ($\beta$ =-7.97, *CI* [-18.67, 2.73], *SE* = 5.51, *t*(190) = -1.45, *P* = 0.15 with exercise-induced pain modulation in the (C) SAL and (D) NLX condition showing no significant difference between males (red) and females (blue). In the NLX condition males show a trend where with increasing fitness levels (FTP) the hypoalgesic response diminished.

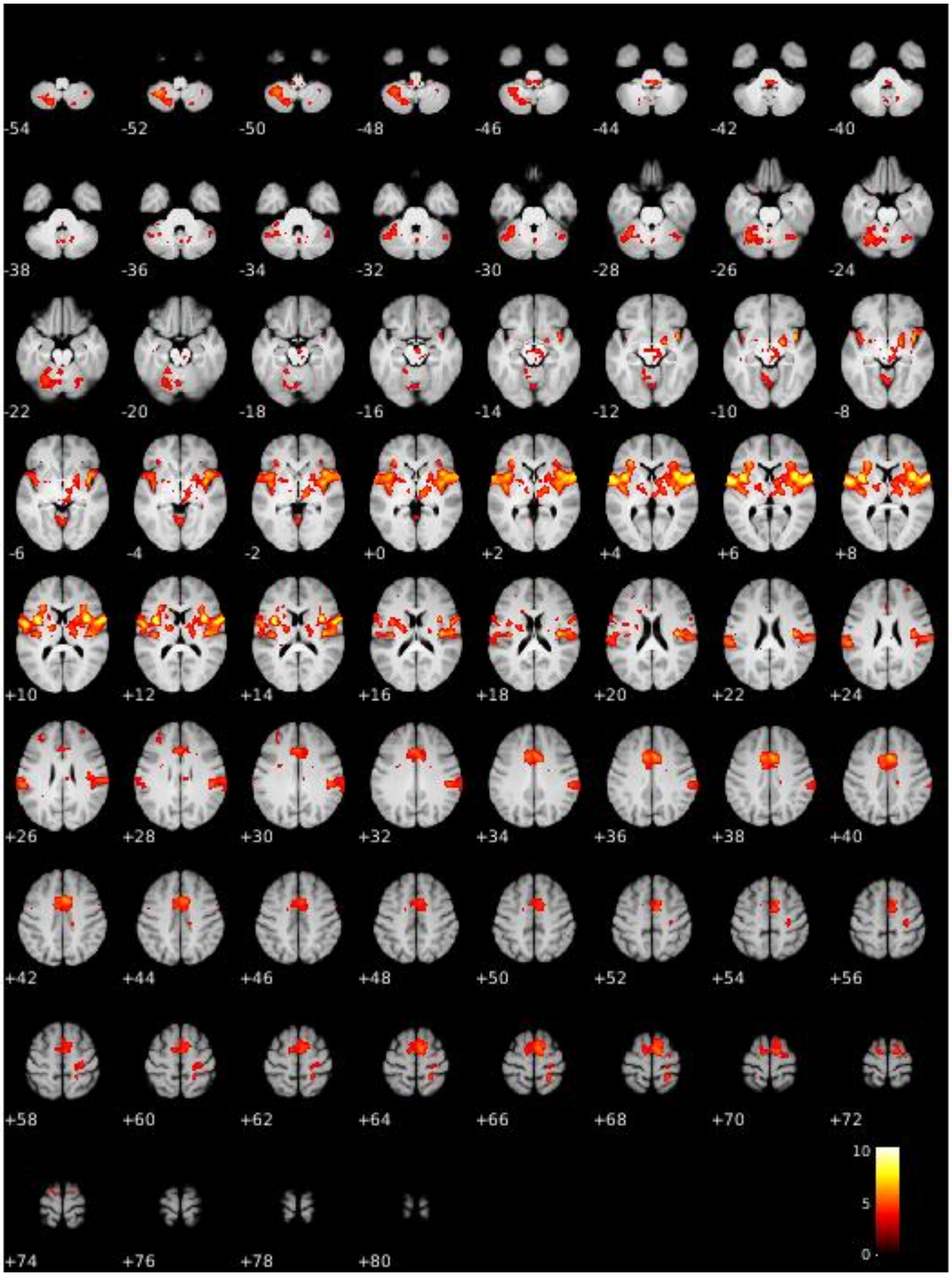

**Figure S2.** Uncorrected activation map for parametric contrast (VAS 70 > 50 > 30) of heat pain in saline (SAL) condition ($P < 0.001$). BOLD activation at $P < 0.001$ uncorrected superimposed onto mean T1 along 134 slices (2 slice steps) in the z-axis.

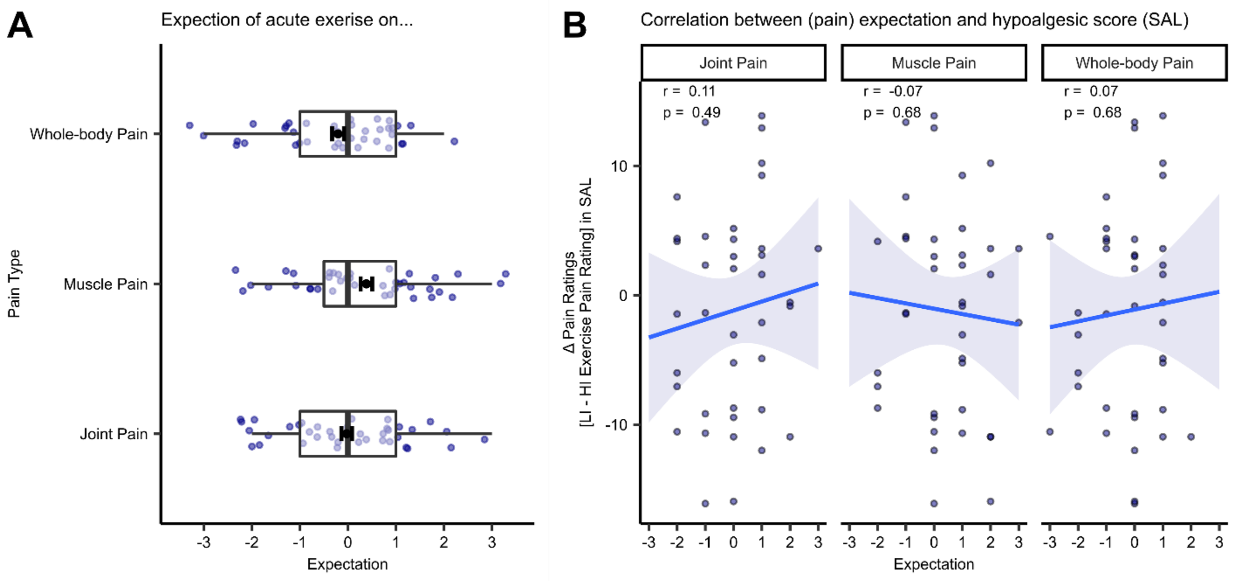

**Figure S3.** Expectation of acute exercise on pain. (A) Ratings for acute exercise on different pain types and ranging between -3 (‘large decrease’) and 3 (‘large increase’) with 0 denoting ‘no effect’. There was no significant effect for muscle pain ($t$(38) = 1.78, $P$ = 0.08, $M$ = 0.39, $SE$ = 0.12), joint pain ($t$(38) = -0.12, $P$ = 0.90, $M$ = -0.03, $SE$ = 0.11), or ‘whole-body pain ($t$(38) = -1.05, $P$ = 0.30, $M$ = -0.21, $SE$ = 0.12) suggesting there to be no expectation effect on these pain domains in the overall sample. Boxplots depict the distribution of the data with dots depicting subject-specific expectation ratings. Black dots depict the mean expectation ratings averaged across subjects and error bars depict the SEM. (B) Correlation between difference pain ratings (LI-HI exercise intensity) in the saline (SAL) condition and expectation ratings with each facet depicting a different pain type for each subject (dot). This analysis yielded no significant correlation in either of the pain domains (joint pain: $r$ = 0.11, $P$ = 0.49; muscle pain: $r$ = -0.07, $P$ = 0.68; whole-body pain: $r$ = 0.07, $P$ = 0.68). Regression lines are visualised and shaded areas represent the SEM ($N$ = 39).

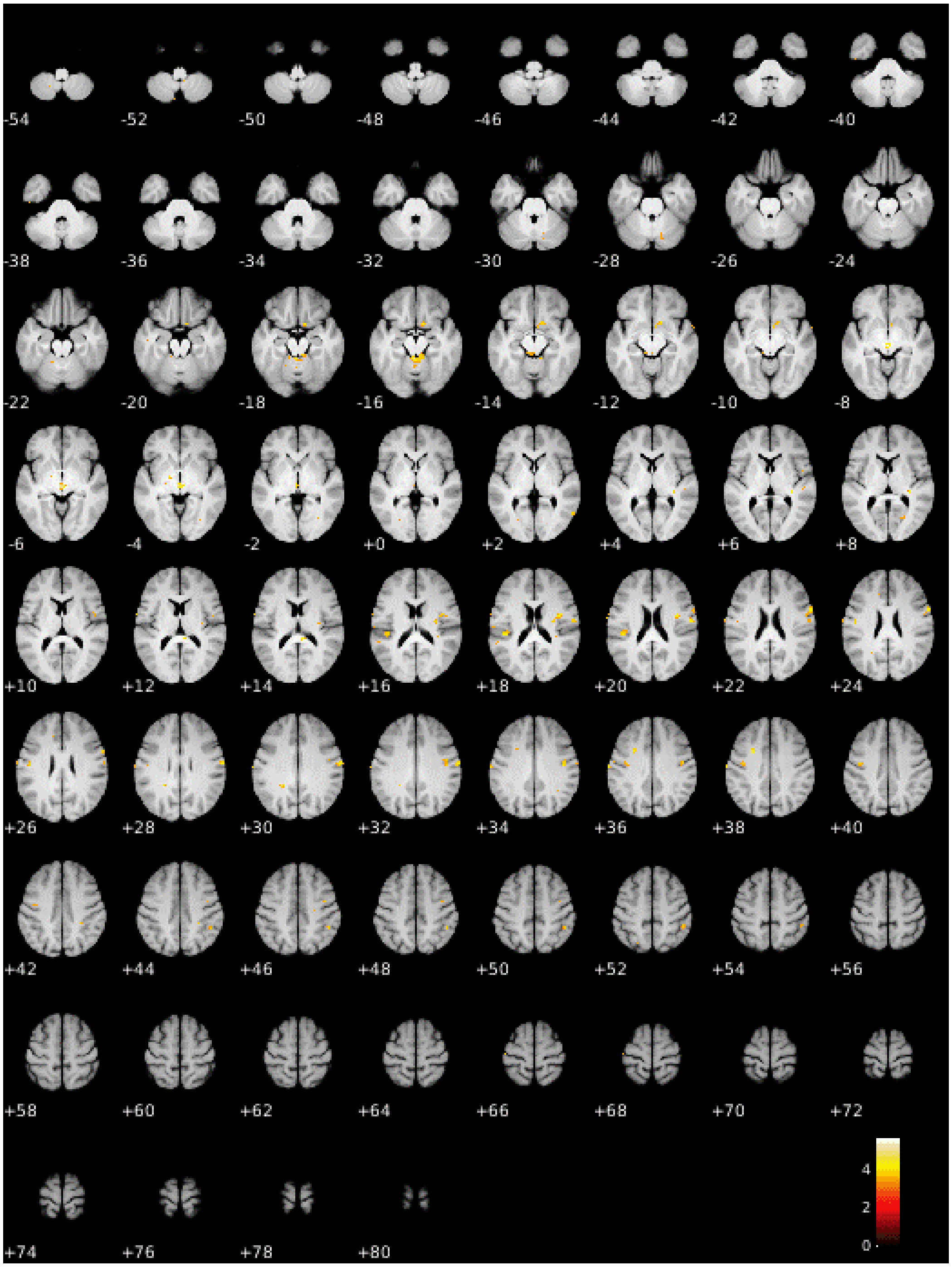

**Figure S4.** Uncorrected activation map for contrast interaction stimulus intensity x drug treatment ($P < 0.001$). BOLD activation at $P < 0.001$ uncorrected superimposed onto mean T1 along 134 slices (2 slice steps) in the z-axis.

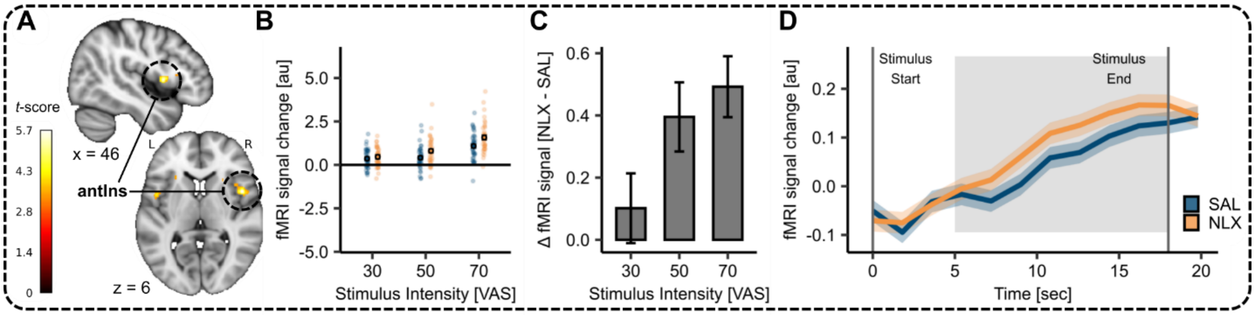

**Figure S5.** Cortical drug treatment effect in the anterior Insula. (**A**) Activation for contrast: heat NLX > heat SAL in the right anterior insula ($MNI_{xyz}$: 46, 8, 6; T = 5.70, $P_{corr\text{-}WB}$ = 0.08) superimposed onto MNI template brain along x and z dimension. (**B**) Parameter estimates for saline (blue) and naloxone (orange) conditions for the respective peak voxel in the right anterior insula ($MNI_{xyz}$: 46, 8, 6). Individual dots indicate subject-specific mean parameter estimates whereas solid dots indicate the overall mean for drug treatment conditions at stimulus intensity. (**C**) Difference between parameter estimates of the respective peak voxels between naloxone and saline condition at each stimulus intensity. (**D**) Time course of BOLD response for saline (blue) and naloxone (orange) condition for heat pain at VAS 70 in the respective peak voxels. The shaded areas around the curves represent the standard error of the mean (*n* = 39). The grey solid lines indicate stimulus start and stimulus end and the shaded grey area displays the approximate time window for BOLD response (5 seconds after stimulus onset).

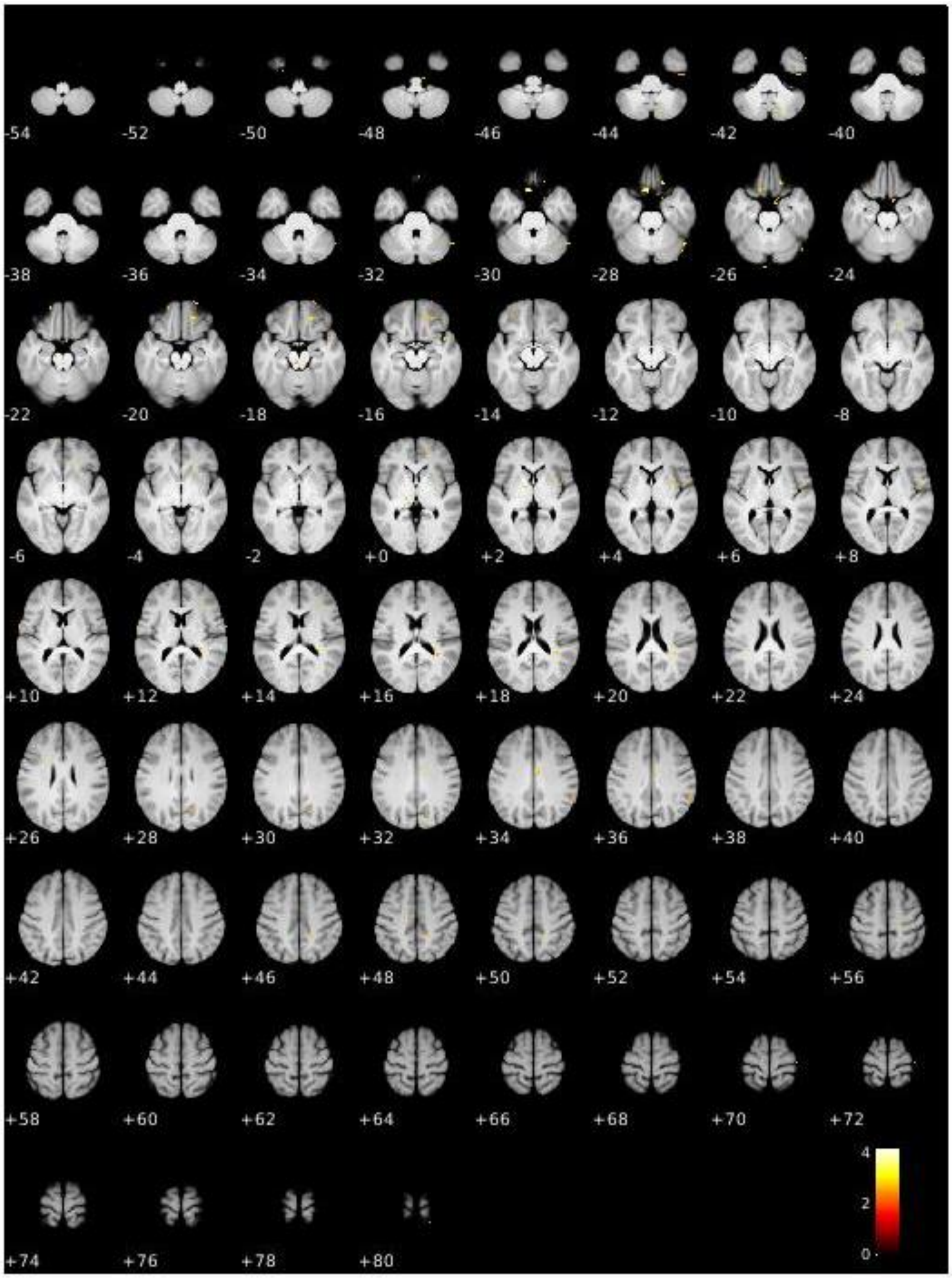

**Figure S6.** Uncorrected activation map for contrast exercise high > low of heat pain in saline (SAL) condition ($P < 0.01$). BOLD activation at $P < 0.01$ uncorrected superimposed onto mean T1 along 134 slices (2 slice steps) in the z-axis.

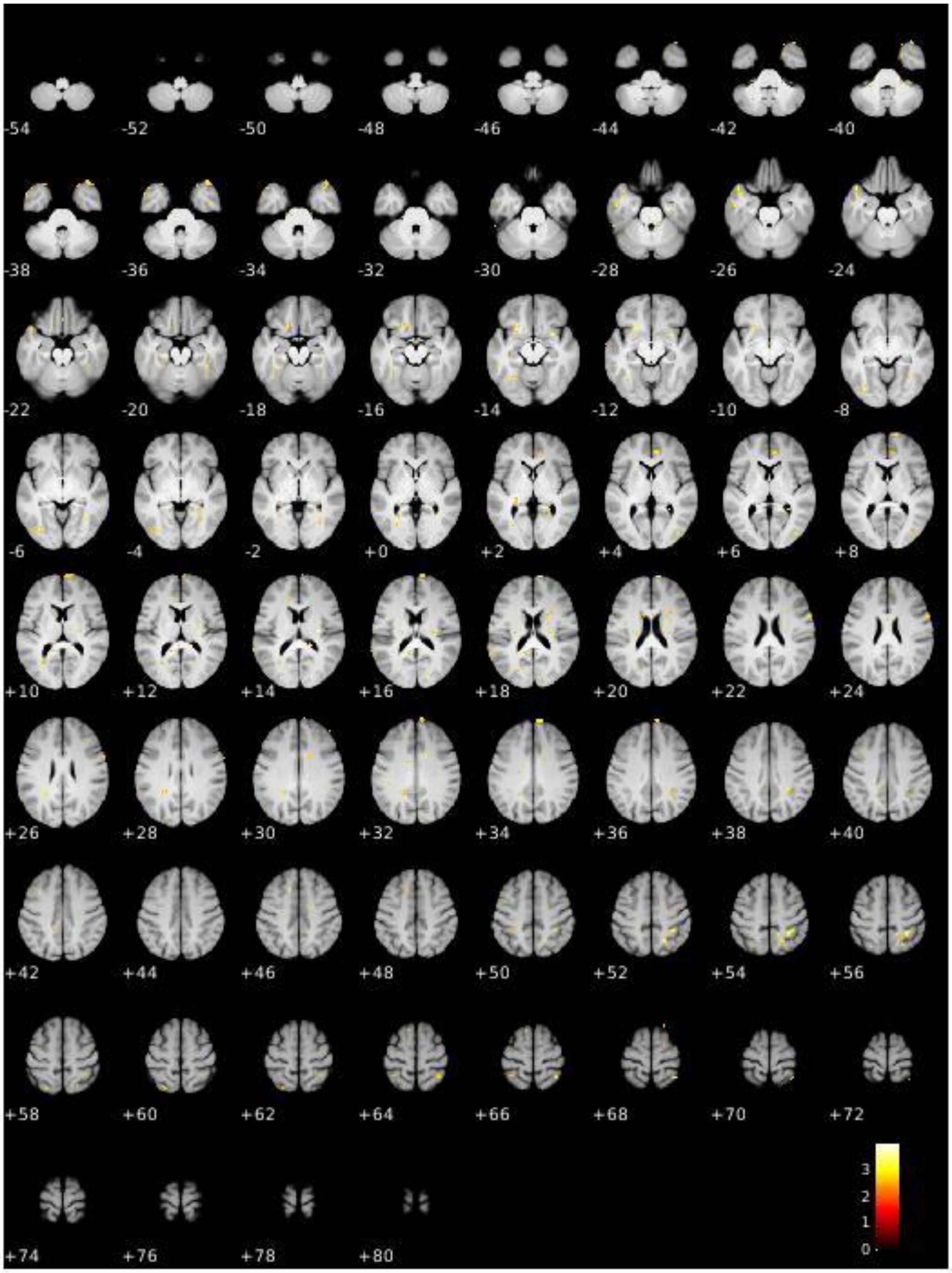

**Figure S7.** Uncorrected activation map for contrast exercise low > high in heat pain in saline (SAL) condition ($P < 0.01$). BOLD activation at $P < 0.01$ uncorrected superimposed onto mean T1 along 134 slices (2 slice steps) in the z-axis.

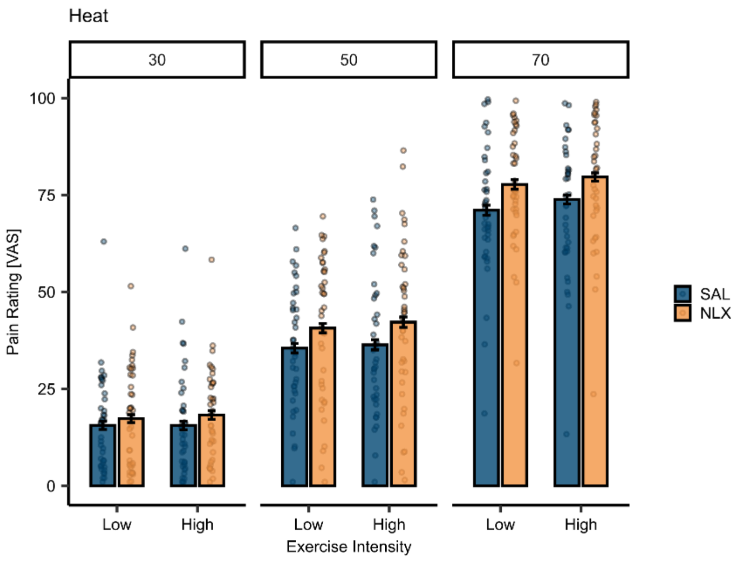

**Figure S8.** Effect of exercise intensity and drug treatment on heat pain ratings at different stimulus intensities (VAS 30, 50, 70). Bars depict the mean ratings in the saline (SAL; blue) and naloxone (NLX; orange) conditions. Individual data points depict subject-specific mean pain ratings. Error bars depict the SEM. The LMER was extended to include the stimulus intensity and yielded a significant main effect of stimulus intensity ($\beta$ = 1.39, *CI* [1.31, 1.47], *SE* = 0.04, $t(2753.12) = -34.082$, $P < 0.001$) and a significant interaction of stimulus intensity and drug treatment ($\beta$ = 0.12, *CI* [0.01, 0.24], *SE* = 0.06, $t(2751) = 2.13$, $P = 0.03$), but no significant interaction of exercise intensity, drug treatment, and stimulus intensity ($\beta$ = -0.05, *CI* [-0.20, 0.11], *SE* = 0.08, $t(2751) = -0.56$, $P = 0.58$).

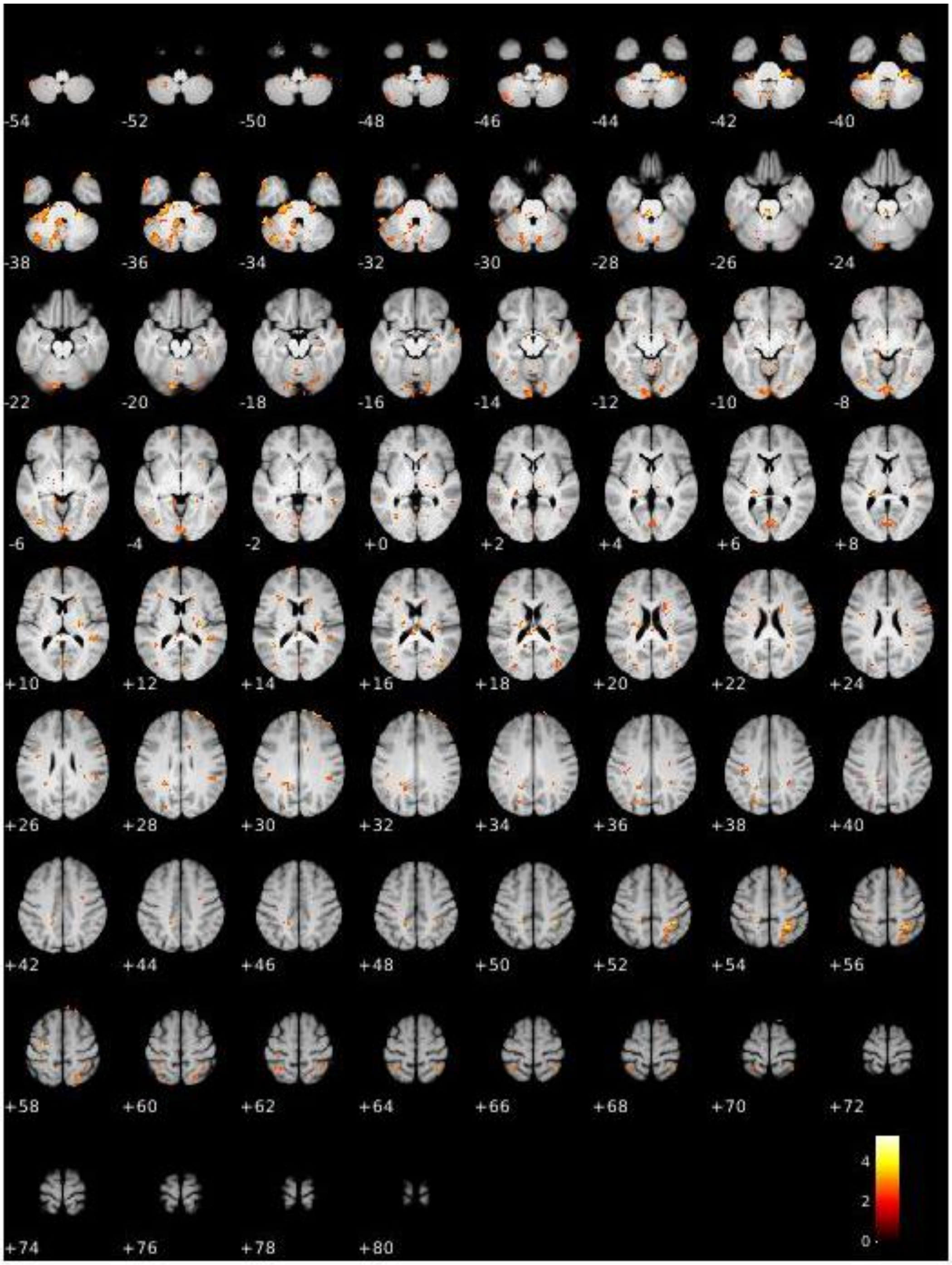

**Figure S9.** Uncorrected activation map for contrast interaction exercise intensity and drug treatment (pos) of heat pain ($P < 0.01$). BOLD activation at $P < 0.01$ uncorrected superimposed onto mean T1 along 134 slices (2 slice steps) in the z-axis.

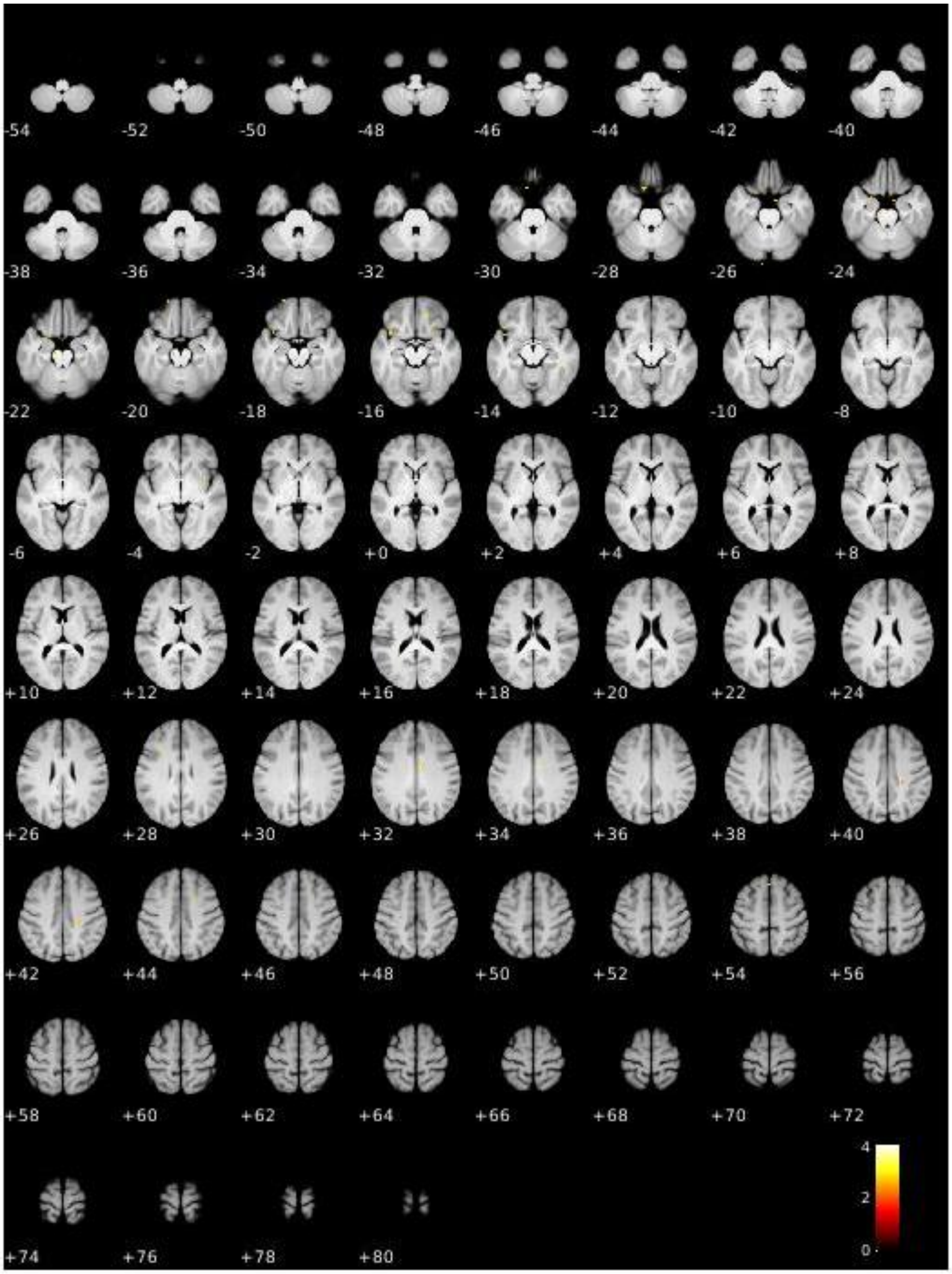

**Figure S10.** Uncorrected activation map for contrast interaction exercise intensity and drug treatment (neg) of heat pain ($P < 0.01$). BOLD activation at $P < 0.01$ uncorrected superimposed onto mean T1 along 134 slices (2 slice steps) in the z-axis.

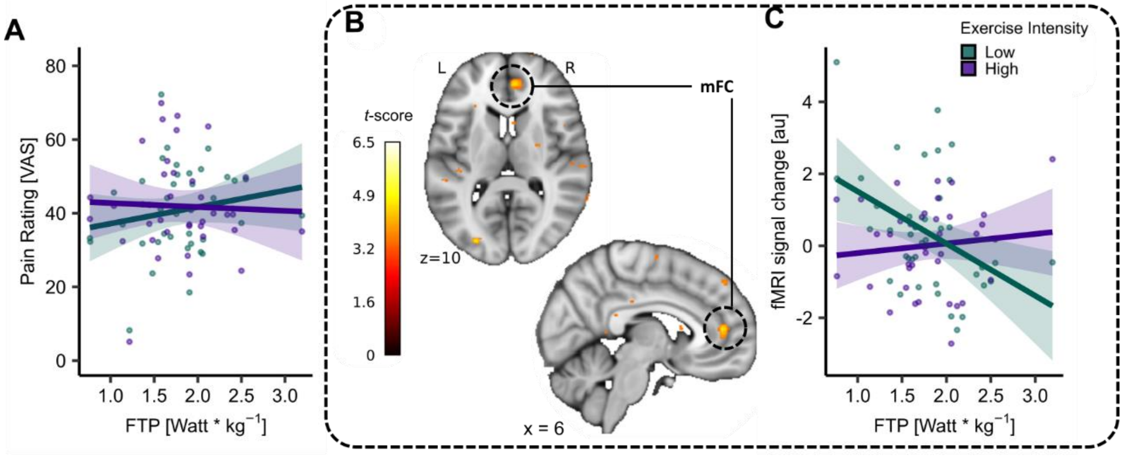

**Figure S11.** Effect of exercise intensity and fitness level on absolute pain ratings and medial frontal cortex (mFC) activation. (**A**) Subject-specific heat pain ratings (dots) between low-intensity (green) and high-intensity (purple) exercise conditions and corresponding regression lines pooled across all stimulus intensities in the saline condition. (**B**) Cortical activation for contrast: exercise high > exercise low with mean-centered covariate FTP (weight-corrected) in right mFC ($MNI_{xyz}$: 6, 45, 10; T = 4.59, $P_{corr\text{-}SVC}$ = 0.05) across all stimulus intensities in the saline condition superimposed onto the MNI template brain. (**C**) Parameter estimates of low-intensity (LI; green) and high-intensity (HI; purple) exercise conditions from respective peak voxel plotted for each subject depending on fitness levels (FTP) and pooled across stimulus intensities.

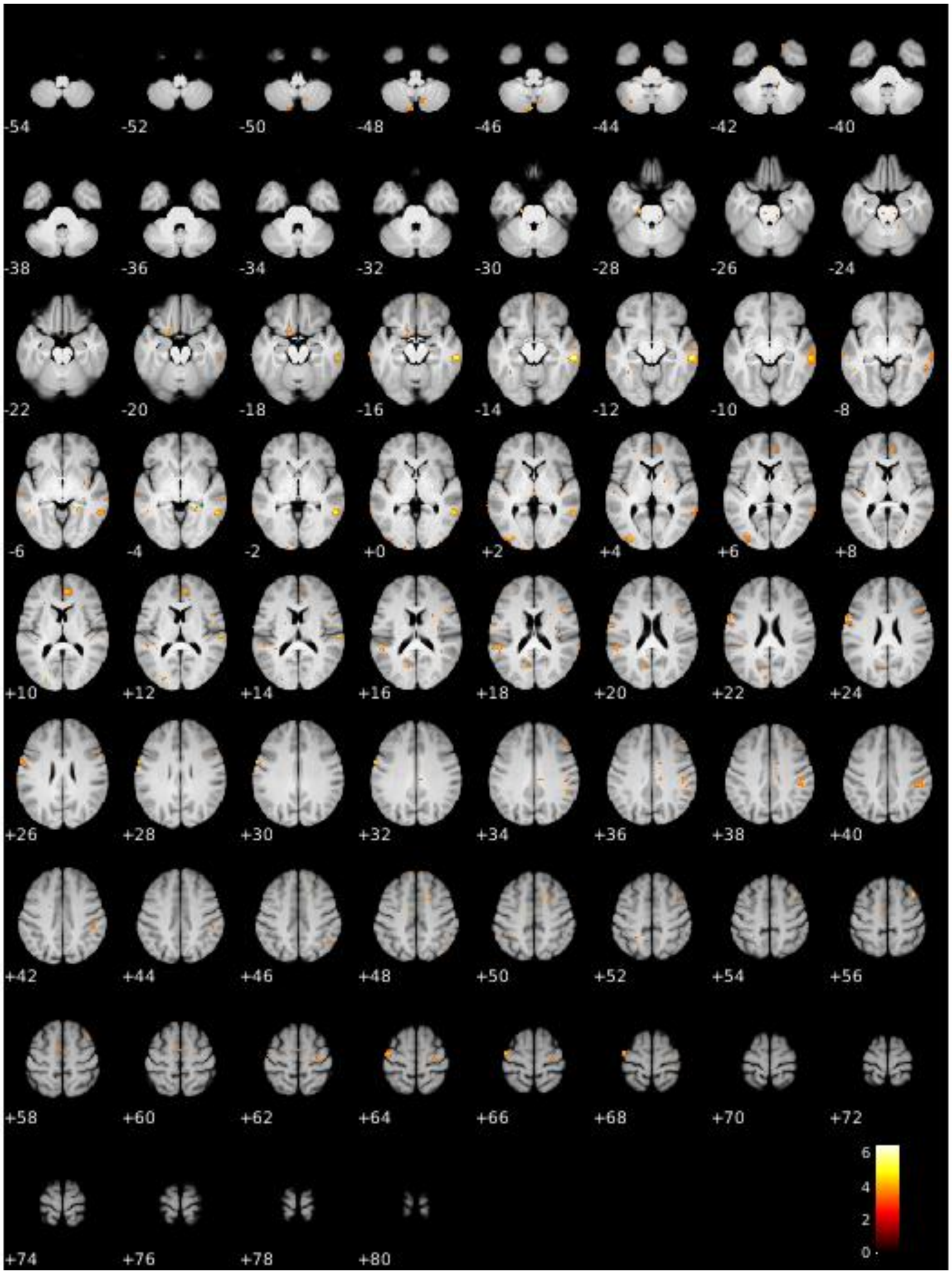

**Figure S12.** Uncorrected activation map for contrast exercise high > exercise low intensity with covariate FTP in saline condition ($P < 0.001$). BOLD activation at $P < 0.001$ uncorrected superimposed onto mean T1 along 134 slices (2 slice steps) in the z-axis.

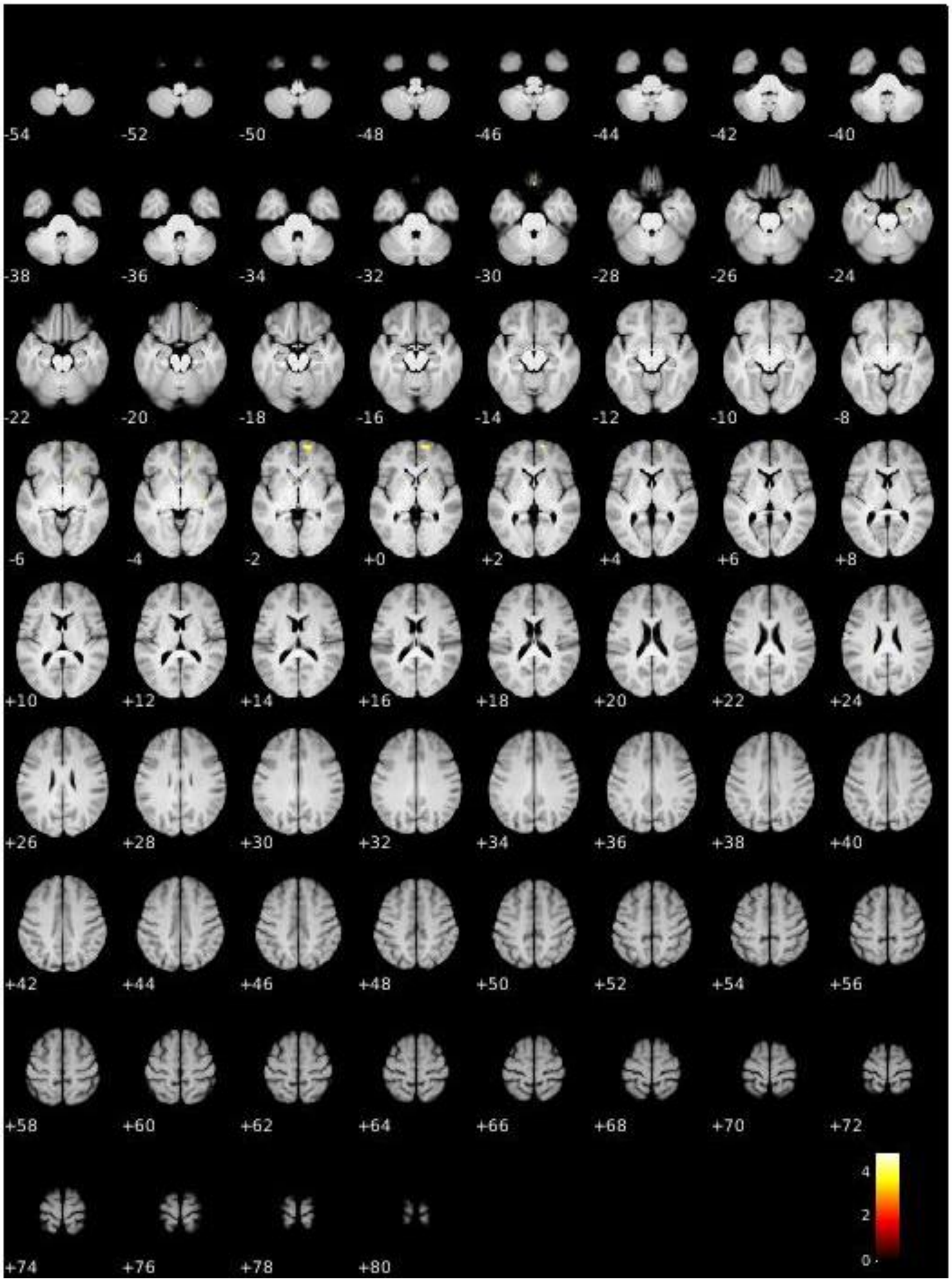

**Figure S13.** Uncorrected activation map for two-sample *t*-test between males and females for contrast: interaction exercise x drug with covariate FTP ($P < 0.001$). BOLD activation at $P < 0.001$ uncorrected superimposed onto mean T1 along 134 slices (2 slice steps) in the z-axis.

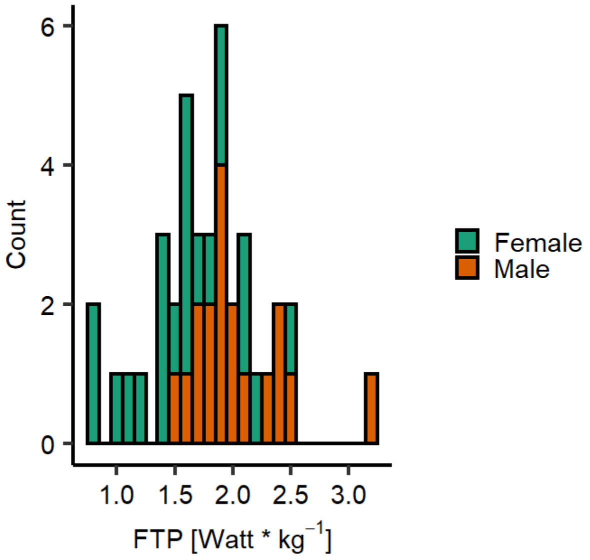

**Figure S14.** Distribution of weight-corrected functional threshold power (FTP) for both sexes. Histogram for distribution of weight-corrected functional threshold power (FTP) for males (orange) and females (green).

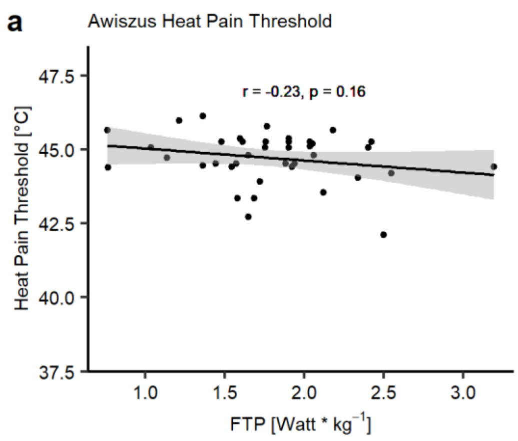

**Figure S15.** Association between heat pain thresholds and fitness level. (A) Heat pain threshold (Awiszus method) as determined during calibration with binary response ('Was this stimulus at least minimally painful?') and fitness level (FTP) as determined by $FTP_{20}$ Test on calibration day show no correlation (*r* = -0.23, *P* = 0.16).

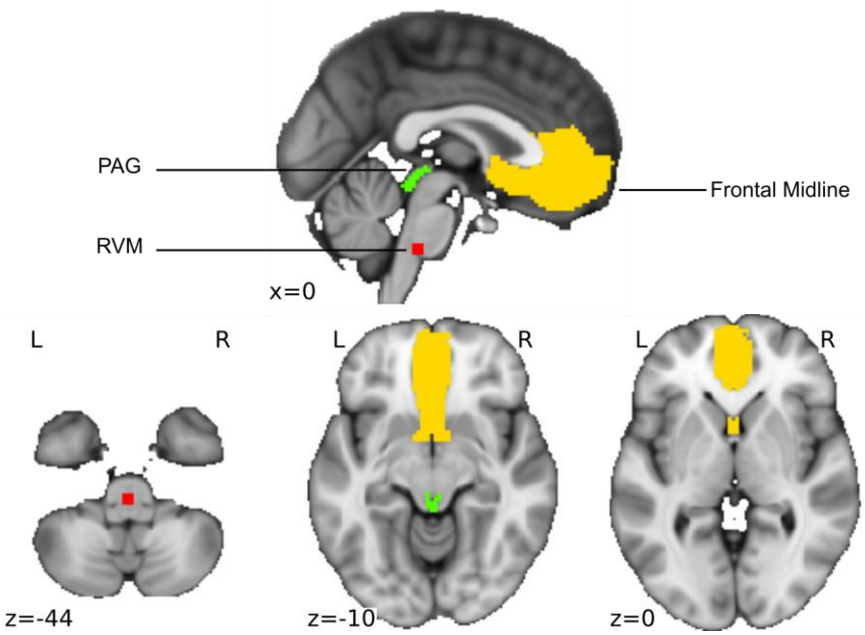

**Figure S16.** Small Volume Correction mask based on preregistered ROIs. Small volume correction mask with 1 mm smoothing in MNI space superimposed onto MNI Template with slices along x (top) and z-axis (bottom). Colour codes are according to ROI: red = rostral ventral medulla (RVM); yellow = Frontal Midline (comprises anterior cingulate cortex (ACC) and ventromedial prefrontal cortex (vmPFC), green = PAG.

**Table S1. Functional Threshold Power Test Protocol.**

| Phase | Duration | Description | Cadence (RPM) | Details |
|---|---|---|---|---|
| Warm-Up | 20-min | Endurance Pace | 65-70 | |
| Intervals | 3 x 1-min | Fast pedalling | 90-100 | 1-min recovery between intervals |
| Recovery | 5-min | Active Recovery | 65 | |
| Time-Trial | 5-min | Maximum Effort | 65-75 | Maximum effort that can be maintained constantly for 5 minutes in a steady state |
| Recovery | 10-min | Active Recovery | 65 | |
| FTP Test | 20-min | Maximum Effort | 65 - 75 | Maximum effort that can be maintained constantly for 20 minutes in a steady state |
| Cool Down | 10-min | Active Recovery | 65 | |

FTP = Functional Threshold Power; RPM = revolutions per minute

**Table S2.** $X^2$**-test to test for differences in the distribution of menstrual cycle phases (*n* = 17 female participants).**

| | $X^2$ | *df* | *P* |
|---|---|---|---|
| Experimental Day 2 | 0.21 | 3 | 0.98 |
| Experimental Day 3 | 0.77 | 3 | 0.86 |
| Exp Day 2 vs. Exp Day 3 | 15.80 | 9 | 0.07 |

$X^2$-test to test for differences in the distribution of menstrual cycle phases (follicular, ovulatory, and luteal phase) for female participants on both experimental days separately and between experimental days (*n* = 17). *N* = 4 female participants were on hormonal contraceptives and were not included in this analysis.*df* = degrees of freedom.

**Table S3. Full LMER model output of parametric effect on behavioural heat pain ratings in saline condition.**

| Fixed Effects | Estimate | *SE* | *df* | *t* | *P* |
|---|---|---|---|---|---|
| Intercept | -34.33 | 2.77 | 70 | -12.40 | $<2\times10^{-16}$ |
| Stimulus Intensity | 1.42 | 0.03 | 1358 | 51.84 | $<2\times10^{-16}$ |
| Treatment_order | 10.22 | 3.51 | 37 | 2.91 | 0.006 |

LMER = linear mixed effects model, *SE* = standard error, *df* = degrees of freedom. Subject and number of pain ratings were included as random effects.

**Table S4. Post-hoc paired *t*-tests (Tukey adj.) for LMER model parametric effect saline behavioural heat pain ratings with according effect size (Cohens *d*).**

| Contrast | Estimate | *SE* | *df* | *t-ratio* | *P* | *Cohens* \|*d*\| |
|---|---|---|---|---|---|---|
| VAS 30 – VAS 50 | -20.4 | 1.07 | 1359 | -18.99 | < 0.001 | 1.25 |
| VAS 30 – VAS 70 | -57.0 | 1.07 | 1359 | -53.24 | < 0.001 | 3.49 |
| VAS 50 – VAS 70 | -36.6 | 1.07 | 1359 | -34.16 | < 0.001 | 2.37 |

LMER = linear mixed effect model, *SE* = standard error, *df* = degrees of freedom

**Table S5. Full LMER model output of stimulus intensity and drug on behavioural heat pain ratings.**

| Fixed Effects | Estimate | *SE* | *df* | *t* | *P* |
|---|---|---|---|---|---|
| Intercept | -31.18 | 2.68 | 80 | -10.89 | $<2\times10^{-16}$ |
| Stimulus Intensity | 1.43 | 0.03 | 2757 | 49.43 | $<2\times10^{-16}$ |
| Drug | -0.37 | 2.14 | 2755 | -0.17 | 0.86 |
| Treatment_order | 2.82 | 3.48 | 37 | 0.81 | 0.42 |
| Stimulus Intensity: drug | 0.10 | 0.04 | 2755 | 2.46 | 0.01 |

LMER = linear mixed effects model, *SE* = standard error, *df* = degrees of freedom. Subject and number of pain ratings were included as random effects.

**Table S6. Post-hoc paired *t*-tests (Tukey adj.) for LMER model interaction stimulus intensity and drug on heat pain ratings.**

| Contrast | Estimate | *SE* | *df* | *t-ratio* | *P* | *Cohens \|d\|* |
|---|---|---|---|---|---|---|
| SAL 30 – NLX 30 | -2.22 | 1.13 | 2753 | -1.97 | 0.36 | 0.13 |
| SAL 30 – SAL 50 | -20.51 | 1.13 | 2755 | -18.10 | < 0.001 | 1.19 |
| SAL 30 – NLX 50 | -25.98 | 1.13 | 2755 | -22.94 | < 0.001 | 1.50 |
| SAL 30 – SAL 70 | -57.04 | 1.13 | 2755 | -50.40 | < 0.001 | 3.30 |
| SAL 30 – NLX 70 | -63.27 | 1.13 | 2755 | -55.91 | < 0.001 | 3.66 |
| NLX 30 – SAL 50 | -18.29 | 1.13 | 2755 | -16.13 | < 0.001 | 1.06 |
| NLX 30 – NLX 50 | -23.76 | 1.13 | 2755 | -20.98 | < 0.001 | 1.38 |
| NLX 30 – SAL 70 | -54.81 | 1.13 | 2755 | -48.44 | < 0.001 | 3.17 |
| NLX 30 – NLX 70 | -61.04 | 1.13 | 2755 | -53.95 | < 0.001 | 3.53 |
| SAL 50 – NLX 50 | -5.47 | 1.13 | 2753 | -4.84 | < 0.001 | 0.32 |
| SAL 50 – SAL 70 | -36.52 | 1.13 | 2755 | -32.24 | < 0.001 | 2.11 |
| SAL 50 – NLX 70 | -42.76 | 1.13 | 2755 | -37.74 | < 0.001 | 2.47 |
| NLX 50 – SAL 70 | -31.05 | 1.13 | 2755 | -27.43 | < 0.001 | 1.80 |
| NLX 50 – NLX 70 | -37.28 | 1.13 | 2755 | -32.93 | < 0.001 | 2.16 |
| SAL 70 – NLX 70 | -6.23 | 1.13 | 2753 | -5.52 | < 0.001 | 0.36 |

LMER = linear mixed effect model, SAL = saline, NLX = naloxone, *SE* = standard error, *df* = degrees of freedom.

**Table S7. Full LMER model output of stimulus intensity on behavioural differential heat pain ratings [NLX – SAL].**

| Fixed Effects | Estimate | *SE* | *df* | *t* | *P* |
|---|---|---|---|---|---|
| Intercept | 6.10 | 2.87 | 100.36 | 2.13 | 0.04 |
| Stimulus Intensity | 0.10 | 0.04 | 77 | 2.46 | 0.02 |
| Treatment_order | -14.80 | 3.07 | 37 | -4.83 | $2.4\times10^{-5}$ |

LMER = linear mixed effects model, *SE* = standard error, *df* = degrees of freedom, NLX = naloxone, SAL = saline. Subject and number of pain ratings were included as random effects.

**Table S8. Post-hoc paired *t*-tests (Tukey adj.) for LMER model interaction stimulus intensity and drug on heat pain ratings for females.**

| Contrast | Estimate | *SE* | *df* | *t-ratio* | *P* | *Cohens \|d\|* |
|---|---|---|---|---|---|---|
| SAL 30 – NLX 30 | 0.16 | 1.57 | 1478 | 0.10 | 1.00 | 0.01 |
| SAL 30 – SAL 50 | -18.71 | 1.57 | 1478 | -11.93 | < 0.001 | 1.06 |
| SAL 30 – NLX 50 | -21.46 | 1.57 | 1478 | -13.67 | < 0.001 | 1.22 |
| SAL 30 – SAL 70 | -53.28 | 1.57 | 1478 | -33.99 | < 0.001 | 3.03 |
| SAL 30 – NLX 70 | -59.93 | 1.57 | 1478 | -38.23 | < 0.001 | 3.41 |
| NLX 30 – SAL 50 | -18.87 | 1.57 | 1478 | -12.03 | < 0.001 | 1.07 |
| NLX 30 – NLX 50 | -21.61 | 1.57 | 1478 | -13.77 | < 0.001 | 1.23 |
| NLX 30 – SAL 70 | -53.44 | 1.57 | 1478 | -34.09 | < 0.001 | 3.04 |
| NLX 30 – NLX 70 | -60.09 | 1.57 | 1478 | -38.34 | < 0.001 | 3.42 |
| SAL 50 – NLX 50 | -2.74 | 1.57 | 1478 | -1.75 | 0.50 | 0.16 |
| SAL 50 – SAL 70 | -34.56 | 1.57 | 1478 | -22.01 | < 0.001 | 1.97 |
| SAL 50 – NLX 70 | -41.22 | 1.57 | 1478 | -26.25 | < 0.001 | 2.35 |
| NLX 50 – SAL 70 | -31.82 | 1.57 | 1478 | -20.27 | < 0.001 | 1.81 |
| NLX 50 – NLX 70 | -38.48 | 1.57 | 1478 | -24.50 | < 0.001 | 2.19 |
| SAL 70 – NLX 70 | -6.66 | 1.57 | 1478 | -4.25 | 0.003 | 0.38 |

LMER = linear mixed effect model, SAL = saline, NLX = naloxone, *SE* = standard error, *df* = degrees of freedom.

**Table S9. Post-hoc paired *t*-tests (Tukey adj.) for LMER model interaction stimulus intensity and drug on heat pain ratings for males.**

| Contrast | Estimate | *SE* | *df* | *t-ratio* | *P* | *Cohens \|d\|* |
|---|---|---|---|---|---|---|
| SAL 30 – NLX 30 | -5.00 | 1.62 | 1262 | -3.10 | 0.02 | 0.03 |
| SAL 30 – SAL 50 | -22.46 | 1.63 | 1262 | -13.79 | < 0.001 | 1.34 |
| SAL 30 – NLX 50 | -31.13 | 1.63 | 1262 | -19.14 | < 0.001 | 1.85 |
| SAL 30 – SAL 70 | -61.34 | 1.62 | 1262 | -37.90 | < 0.001 | 3.65 |
| SAL 30 – NLX 70 | -67.08 | 1.62 | 1262 | -41.45 | < 0.001 | 4.00 |
| NLX 30 – SAL 50 | -17.45 | 1.63 | 1262 | -10.72 | < 0.001 | 1.04 |
| NLX 30 – NLX 50 | -26.13 | 1.63 | 1262 | -16.06 | < 0.001 | 1.56 |
| NLX 30 – SAL 70 | -56.33 | 1.62 | 1262 | -34.81 | < 0.001 | 3.36 |
| NLX 30 – NLX 70 | -62.07 | 1.62 | 1262 | -38.35 | < 0.001 | 3.70 |
| SAL 50 – NLX 50 | -8.68 | 1.62 | 1262 | -5.35 | < 0.001 | 0.52 |
| SAL 50 – SAL 70 | -38.88 | 1.62 | 1262 | -23.94 | < 0.001 | 2.32 |
| SAL 50 – NLX 70 | -44.62 | 1.62 | 1262 | -27.47 | < 0.001 | 2.66 |
| NLX 50 – SAL 70 | -30.20 | 1.62 | 1262 | -18.62 | < 0.001 | 1.80 |
| NLX 50 – NLX 70 | -35.94 | 1.62 | 1262 | -22.16 | < 0.001 | 2.14 |
| SAL 70 – NLX 70 | -5.74 | 1.62 | 1262 | -3.55 | 0.005 | 0.34 |

LMER = linear mixed effect model, SAL = saline, NLX = naloxone, *SE* = standard error, *df* = degrees of freedom.

**Table S10. Full LMER model output of stimulus intensity and sex on behavioural differential heat pain ratings [NLX – SAL].**

| Fixed Effects | Estimate | *SE* | *df* | *t* | *P* |
|---|---|---|---|---|---|
| Intercept | 0.93 | 3.66 | 104.79 | 0.25 | 0.80 |
| Stimulus Intensity | 0.17 | 0.05 | 76 | 3.12 | 0.003 |
| Sex | 11.26 | 5.03 | 111.12 | 2.24 | 0.03 |
| Treatment_order | -14.85 | 3.05 | 36 | -4.88 | $2.19\times10^{-5}$ |
| Stimulus Intensity: Sex | -0.15 | 0.08 | 76 | -1.89 | 0.06 |

LMER = linear mixed effects model, *SE* = standard error, *df* = degrees of freedom, NLX = naloxone, SAL = saline. Subject and number of pain ratings were included as random effects.

**Table S11. Full LMER output of exercise intensity on heat pain ratings in the saline condition.**

| Fixed Effects | Estimate | *SE* | *df* | *t* | *P* |
|---|---|---|---|---|---|
| Intercept | 36.29 | 2.46 | 53.75 | 14.75 | $2\times10^{-16}$ |
| Exercise Intensity | 1.19 | 1.55 | 1354 | 0.77 | 0.44 |
| Treatment_order | 10.244 | 3.51 | 37 | 2.91 | 0.006 |

LMER = linear mixed effects model, *SE* = standard error, *df* = degrees of freedom. Subject and number of pain ratings were included as random effects.

**Table S12. Full LMER output of exercise intensity on betas extracted from ROI RVM in the saline condition.**

| Fixed Effects | Estimate | *SE* | *df* | *t* | *P* |
|---|---|---|---|---|---|
| Intercept | 0.07 | 0.10 | 49.73 | 0.72 | 0.48 |
| Exercise Intensity | 0.05 | 0.07 | 194 | 0.63 | 0.53 |
| Treatment_order | 0.23 | 0.12 | 37 | 1.96 | 0.06 |

LMER = linear mixed effects model, *SE* = standard error, *df* = degrees of freedom. Subject was included as a random effect.

**Table S13. Full LMER output of exercise intensity on betas extracted from ROI PAG in the saline condition.**

| Fixed Effects | Estimate | *SE* | *df* | *t* | *P* |
|---|---|---|---|---|---|
| Intercept | 0.09 | 0.09 | 51.62 | 0.93 | 0.36 |
| Exercise Intensity | 0.02 | 0.08 | 194 | 0.31 | 0.76 |
| Treatment_order | 0.11 | 0.12 | 37 | 0.96 | 0.35 |

LMER = linear mixed effects model, *SE* = standard error, *df* = degrees of freedom. Subject was included as a random effect.

**Table S14. Full LMER output of exercise intensity on betas extracted from ROI Frontal Midline in the saline condition.**

| Fixed Effects | Estimate | *SE* | *df* | *t* | *P* |
|---|---|---|---|---|---|
| Intercept | -0.16 | 0.09 | 46.61 | -1.67 | 0.10 |
| Exercise Intensity | -0.08 | 0.06 | 194 | -1.26 | 0.21 |
| Treatment_order | 0.19 | 0.12 | 37 | 1.65 | 0.11 |

LMER = linear mixed effects model, *SE* = standard error, *df* = degrees of freedom. Subject was included as a random effect.

**Table S15. Full LMER output of exercise intensity and drug treatment on heat pain ratings.**

| Fixed Effects | Estimate | *SE* | *df* | *t* | *P* |
|---|---|---|---|---|---|
| Intercept | 36.52 | 3.00 | 54.3 | 15.23 | $2\times10^{-16}$ |
| Exercise Intensity | 1.19 | 1.60 | 2755 | 0.75 | 0.45 |
| Drug | 4.50 | 1.60 | 2755 | 2.81 | 0.005 |
| Treatment_order | 2.82 | 3.48 | 37 | 0.81 | 0.42 |
| Exercise Intensity x Drug | 0.27 | 2.27 | 2755 | 0.12 | 0.91 |

LMER = linear mixed effects model, *SE* = standard error, *df* = degrees of freedom. Subject and number of pain ratings were included as random effects.

**Table S16. Full LMER output of exercise intensity and drug treatment on betas extracted from ROI RVM.**

| Fixed Effects | Estimate | *SE* | *df* | *t* | *P* |
|---|---|---|---|---|---|
| Intercept | 0.04 | 0.09 | 44.63 | 0.49 | 0.63 |
| Exercise Intensity | 0.06 | 0.05 | 426 | 1.05 | 0.29 |
| Drug | -0.02 | 0.04 | 426 | -0.15 | 0.68 |
| Treatment_order | 0.25 | 0.12 | 37 | 2.15 | 0.04 |
| Exercise Intensity x Drug | 0.01 | 0.05 | 426 | 0.23 | 0.82 |

LMER = linear mixed effects model, *SE* = standard error, *df* = degrees of freedom. Subject was included as a random effect.

**Table S17. Full LMER output of exercise intensity and drug treatment on betas extracted from ROI PAG.**

| Fixed Effects | Estimate | *SE* | *df* | *t* | *P* |
|---|---|---|---|---|---|
| Intercept | 0.13 | 0.08 | 48.45 | 1.68 | 0.10 |
| Exercise Intensity | 0.08 | 0.06 | 426 | 1.42 | 0.16 |
| Drug | 0.05 | 0.04 | 426 | 1.17 | 0.24 |
| Treatment_order | 0.11 | 0.10 | 37 | 1.18 | 0.25 |
| Exercise Intensity x Drug | 0.06 | 0.06 | 426 | 1.00 | 0.32 |

LMER = linear mixed effects model, *SE* = standard error, *df* = degrees of freedom. Subject was included as a random effect.

**Table S18. Full LMER output of exercise intensity and drug treatment on betas extracted from ROI frontal midline.**

| Fixed Effects | Estimate | *SE* | *df* | *t* | *P* |
|---|---|---|---|---|---|
| Intercept | -0.15 | 0.07 | 46.44 | -2.16 | 0.04 |
| Exercise Intensity | -0.05 | 0.04 | 426 | -1.12 | 0.26 |
| Drug | -0.03 | 0.03 | 426 | .1.09 | 0.28 |
| Treatment_order | 0.12 | 0.09 | 37 | 1.36 | 0.18 |
| Exercise Intensity x Drug | 0.03 | 0.04 | 426 | 0.63 | 0.53 |

LMER = linear mixed effects model, *SE* = standard error, *df* = degrees of freedom. Subject was included as a random effect.

**Table S19. Full linear model output from the model including FTP on difference score heat pain ratings (LI – HI exercise) in the saline condition.**

| Fixed Effects | Estimate | *SE* | *t* | *P* |
|---|---|---|---|---|
| Intercept | -10.84 | 4.66 | -2.33 | 0.03 |
| FTP | 6.45 | 2.56 | 2.52 | 0.02 |
| Treatment_order | -4.33 | 2.49 | -1.74 | 0.09 |

FTP = functional threshold power (weight-corrected), *SE* = standard error, *df* = degrees of freedom. Subject and number of pain ratings were included as random effects.

**Table S20. LMER output from the model including FTP, drug, and sex as fixed effects on differential heat pain ratings (LI exercise – HI exercise).**

| Fixed Effects | Estimate | *SE* | *df* | *t* | *P* |
|---|---|---|---|---|---|
| intercept | -1.52 | 6.09 | 67.27 | -0.25 | 0.80 |
| Stimulus intensity | -0.05 | 0.03 | 190 | -1.35 | 0.18 |
| FTP | 2.62 | 3.61 | 56.77 | 0.73 | 0.47 |
| Sex | -24.28 | 10.56 | 57 | -2.30 | 0.03 |
| Drug | -9.76 | 5.72 | 190 | -1.71 | 0.09 |
| Treatment_order | -4.72 | 2.09 | 34 | -2.26 | 0.03 |
| Sex: Drug | 28.61 | 10.34 | 190 | 2.77 | 0.006 |
| FTP: Drug | 4.81 | 3.48 | 190 | 1.39 | 0.17 |
| FTP:Sex | 11.98 | 5.54 | 57.61 | 2.16 | 0.03 |
| FTP:Sex: Drug | -13.12 | 5.41 | 190 | -2.43 | 0.016 |

LMER = linear mixed effects model, FTP = functional threshold power (weight-corrected), *SE* = standard error, *df* = degrees of freedom. The subject was included as a random effect.

**Table S21. Participant characteristics.**

| | Overall Mean (SD) | Females Mean (SD) | Males Mean (SD) |
|---|---|---|---|
| *N* | 39 | 21 | 18 |
| Age (years) | 26.03 (4.83) | 25.33 (5.10) | 26.83 (4.35) |
| Weight (kg) | 70.95 (12.14) | 63.33 (7.53) | 79.83 (10.38) |
| Height (cm) | 177.10 (9.08) | 170.52 (6.35) | 184.78 (4.58) |
| BMI | 22.51 (2.64) | 21.80 (2.59) | 23.33 (2.53) |
| FTP (Watt/kg) | 1.79 (0.48) | 1.58 (0.44) | 2.03 (0.40) |
| Training Volume (h/w) | 4.44 (3.24) | 3.88 (2.63) | 5.08 (3.81) |

SD = Standard deviation. kg = kilogram. cm = centimeter. h/w = hours per week.

**Table S22. POMS Mood Ratings (Wilcoxon signed-rank test).**

| | *W* | *P* |
|---|---|---|
| Naloxone Pre - Post | | |
| Fatigue | 544.5 | < 0.001 |
| Drive | 174 | 0.01 |
| Discontent | 22 | 0.05 |
| Dejection | 10.5 | 0.001 |
| Saline Pre - Post | | |
| Fatigue | 381.5 | 0.28 |
| Drive | 409.5 | 0.39 |
| Discontent | 12 | 0.23 |
| Dejection | 33.5 | 0.007 |
| Saline Post – Naloxone Post | | |
| Fatigue | 226 | 0.15 |
| Drive | 437 | 0.19 |
| Discontent | 17 | 0.67 |
| Dejection | 57.5 | 0.42 |

*W* = test statistic.

**Table S23. Side Effects Naloxone (Wilcoxon signed-rank test).**

| | *W* | *P* |
|---|---|---|
| Lethargy | 91 | 0.61 |
| Dry Mouth | 77.5 | 0.06 |
| Dry Skin | 68 | 0.45 |
| Blurred Vision | 25.5 | 0.29 |
| Dizziness | 26 | 0.37 |
| Headache | 6 | 0.37 |
| Sickness | 4 | 0.41 |

*W* = test statistic.

**Table S24. Small Volume Correction (SVC) mask for pain modulation effects based on preregistered midbrain ROIS.**

| Label | Description | Atlas/Source | x | y | z | Sphere (mm) |
|---|---|---|---|---|---|---|
| PAG | Periaqueductal grey | Brainstem Navigator (Bianciardi et al., 2018, 2015; García-Gomar et al., 2022, 2019; Singh et al., 2021, 2020) | - | - | - | - |
| RVM | Rostral ventral medulla | ROI from Tinnermann et al. (2017) (Tinnermann et al., 2017) | 0 | -32 | -44 | 3 |
| Frontal Midline | | | | | | |
| | pgACC bilateral | From Vega et al., (2016) (Vega et al., 2016) | - | - | - | - |
| | vmPFC | From Vega et al., (2016) (Vega et al., 2016) | - | - | - | - |

xyz-coordinates are provided in the MNI space. mm = millimetre, PAG = Periaqueductal gray, pgACC = pregenual anterior cingulate cortex, RVM = Rostral ventral medulla, vmPFC = ventromedial prefrontal cortex.

### Supplemental References

Bianciardi M, Strong C, Toschi N, Edlow BL, Fischl B, Brown EN, Rosen BR, Wald LL. 2018. A probabilistic template of human mesopontine tegmental nuclei from in vivo 7 T MRI. *NeuroImage* **170**:222–230. doi:10.1016/j.neuroimage.2017.04.070

Bianciardi M, Toschi N, Edlow BL, Eichner C, Setsompop K, Polimeni JR, Brown EN, Kinney HC, Rosen BR, Wald LL. 2015. Toward an In Vivo Neuroimaging Template of Human Brainstem Nuclei of the Ascending Arousal, Autonomic, and Motor Systems. *Brain Connect* **5**:597–607. doi:10.1089/brain.2015.0347

García-Gomar MG, Strong C, Toschi N, Singh K, Rosen BR, Wald LL, Bianciardi M. 2019. In vivo Probabilistic Structural Atlas of the Inferior and Superior Colliculi, Medial and Lateral Geniculate Nuclei and Superior Olivary Complex in Humans Based on 7 Tesla MRI. *Front Neurosci* **13**:764. doi:10.3389/fnins.2019.00764

García-Gomar MG, Videnovic A, Singh K, Stauder M, Lewis LD, Wald LL, Rosen BR, Bianciardi M. 2022. Disruption of Brainstem Structural Connectivity in REM Sleep Behavior Disorder Using 7 Tesla Magnetic Resonance Imaging. *Mov Disord* **37**:847–853. doi:10.1002/mds.28895

Singh K, García-Gomar MG, Bianciardi M. 2021. Probabilistic Atlas of the Mesencephalic Reticular Formation, Isthmic Reticular Formation, Microcellular Tegmental Nucleus, Ventral Tegmental Area Nucleus Complex, and Caudal-Rostral Linear Raphe Nucleus Complex in Living Humans from 7 Tesla Magnetic Resonance Imaging. *Brain Connect* **11**:613–623. doi:10.1089/brain.2020.0975

Singh K, Indovina I, Augustinack JC, Nestor K, García-Gomar MG, Staab JP, Bianciardi M. 2020. Probabilistic Template of the Lateral Parabrachial Nucleus, Medial Parabrachial Nucleus, Vestibular Nuclei Complex, and Medullary Viscero-Sensory-Motor Nuclei Complex in Living Humans From 7 Tesla MRI. *Front Neurosci* **13**:1425. doi:10.3389/fnins.2019.01425

Tinnermann A, Geuter S, Sprenger C, Finsterbusch J, Büchel C. 2017. Interactions between brain and spinal cord mediate value effects in nocebo hyperalgesia. *Science* **358**:105–108. doi:10.1126/science.aan1221

Vega A de la, Chang LJ, Banich MT, Wager TD, Yarkoni T. 2016. Large-Scale Meta-Analysis of Human Medial Frontal Cortex Reveals Tripartite Functional Organization. *J Neurosci* **36**:6553–6562. doi:10.1523/JNEUROSCI.4402-15.2016

Wilcoxon F. 1945. Individual Comparisons by Ranking Methods. *Biom Bull* **1**:80–83. doi:10.2307/3001968
